# Homeostatic feedback modulates the development of two-state patterned activity in a model serotonin motor circuit in *Caenorhabditis elegans*

**DOI:** 10.1101/202507

**Authors:** Bhavya Ravi, Jessica Garcia, Kevin M. Collins

## Abstract

Neuron activity accompanies synapse formation and maintenance, but how early circuit activity contributes to behavior development is not well understood. Here, we use the *Caenorhabditis elegans* egg-laying motor circuit as a model to understand how coordinated cell and circuit activity develops and drives a robust two-state behavior in adults. Using calcium imaging in behaving animals, we find the serotonergic Hermaphrodite Specific Neurons (HSNs) and vulval muscles show rhythmic calcium transients in L4 larvae before eggs are produced. HSN activity in L4 is tonic and lacks the alternating burst-firing/quiescent pattern seen in egg-laying adults. Vulval muscle activity in L4 is initially uncoordinated but becomes synchronous as the anterior and posterior muscle arms meet at HSN synaptic release sites. However, coordinated muscle activity does not require presynaptic HSN input. Using reversible silencing experiments, we show that neuronal and vulval muscle activity in L4 is not required for the onset of adult behavior. Instead, the accumulation of eggs in the adult uterus renders the muscles sensitive to HSN input. Sterilization or acute electrical silencing of the vulval muscles inhibits presynaptic HSN activity, and reversal of muscle silencing triggers a homeostatic increase in HSN activity and egg release that maintains ~12-15 eggs in the uterus. Feedback of egg accumulation depends upon the vulval muscle postsynaptic terminus, suggesting a retrograde signal sustains HSN synaptic activity and egg release. Our results show that egg-laying behavior in *C. elegans* is driven by a homeostat that scales serotonin motor neuron activity in response to postsynaptic muscle feedback.

**Significance Statement:** The functional importance of early, spontaneous neuron activity in synapse and circuit development is not well understood. Here we show that in the nematode *C. elegans*, the serotonergic Hermaphrodite Specific Neurons (HSNs) and postsynaptic vulval muscles show activity during circuit development, well before the onset of adult behavior. Surprisingly, early activity is not required for circuit development or the onset of adult behavior, and the circuit remains unable to drive egg laying until fertilized embryos are deposited into the uterus. Egg accumulation potentiates vulval muscle excitability, but ultimately acts to promote burst firing in the presynaptic HSNs during which eggs are laid. Our results suggest that mechanosensory feedback acts at three distinct steps to initiate, sustain, and terminate *C. elegans* egg-laying circuit activity and behavior.

## Introduction

Developing neural circuits in the cortex, hippocampus, cerebellum, retina, and spinal cord show spontaneous neural activity (Wong et al., 1995; Garaschuk et al., 1998; Garaschuk et al., 2000; Watt et al., 2009; Warp et al., 2012). In contrast, mature neural circuits show coordinated patterns of activity required to drive efficient behaviors. Activity-dependent mechanisms have been shown to play key roles during development in vertebrate neural circuits (Gu et al., 1994; Gu and Spitzer, 1995; Jarecki and Keshishian, 1995; Borodinsky et al., 2004; Hanson et al., 2008), but the complexity of such circuits poses limitations in terms of understanding how developmental events, neurotransmitter specification, and sensory signals act together to promote the transition from immature to mature patterns of circuit activity. Genetically tractable invertebrate model organisms, such as the nematode *Caenorhabditis elegans*, have simple neural circuits and are amenable to powerful experimental approaches allowing us to investigate how activity in neural circuits is shaped during development.

The *C. elegans* egg laying circuit is a well-characterized neural circuit that drives a two-state behavior in adult animals with ~20 minute inactive periods punctuated by ~2 minute active states where ~4-6 eggs are laid (Waggoner et al., 1998). The egg-laying circuit comprises two serotonergic Hermaphrodite Specific Neurons (HSNs) which promote the active state (Waggoner et al., 1998; Emtage et al., 2012), three locomotion motor neurons (VA7, VB6, and VD7), and six cholinergic Ventral C neurons (VC1-6), all of whom synapse onto a set of vulval muscles whose rhythmic activity drives either weak twitching or the release of eggs from the uterus in phase with locomotion (White et al., 1986; Collins and Koelle, 2013; Collins et al., 2016). Four uv1 neuroendocrine cells connect the vulval canal to the uterus which holds embryos until they are laid. HSN, VC, uv1, and vulval muscle development occurs during the early-mid L4 larval stages and requires interactions with the developing vulval epithelium, but not the other cells in the circuit (Colavita and Tessier-Lavigne, 2003; Shen and Bargmann, 2003; Shen et al., 2004).

During egg laying, serotonin released from the HSNs signals through vulval muscle receptors (Carnell et al., 2005; Dempsey et al., 2005; Hobson et al., 2006; Dernovici et al., 2007; Hapiak et al., 2009), likely increasing the excitability of the muscles so that rhythmic input from cholinergic motor neurons can drive vulval muscle contractions (White et al., 1986; Collins and Koelle, 2013; Collins et al., 2016). We have previously shown that HSN Ca^2+^ transients occur more frequently during the active state, but the factors which promote this timely ‘feed-forward’ increase in HSN activity remain poorly understood. The cholinergic VCs show rhythmic Ca^2+^ transients coincident with vulval muscle contractions during the active state, although whether VC activity drives contraction itself or instead acts to modulate HSN signaling is still not clear (Bany et al., 2003; Zhang et al., 2008; Zang et al., 2017). The uv1 cells, mechanically deformed by the passage of eggs through the vulva, release tyramine and neuropeptides that signal extrasynaptically to inhibit HSN activity (Collins et al., 2016; Banerjee et al., 2017). Because each cell in the circuit develops independently in juveniles, how this circuit goes on to develop the robust pattern of coordinated activity seen in adults remains unclear.

We show here the presynaptic HSN motor neurons and the postsynaptic vulval muscles are active during the late L4 larval stage, well before egg production and the onset of adult egg-laying behavior. We do not observe activity in the VC neurons and uv1 neuroendocrine cells until behavioral onset. The adult circuit remains in a nonfunctional state until receiving feedback of eggs in the uterus. This egg-laying homeostat requires the vm2 muscle arms and muscle activity which we show promotes HSN burst firing that maintains the active state. Together, our data reveal how cell activity patterns that emerge during circuit development are modulated by sensory feedback that decide when and for how long to drive behavior.

## Materials and Methods

### Nematode Culture and Developmental Staging

*Caenorhabditis elegans* hermaphrodites were maintained at 20°C on Nematode Growth Medium (NGM) agar plates with *E. coli* OP50 as a source of food as described (Brenner, 1974). Animals were staged and categorized based on the morphology of the vulva as described (Mok et al., 2015). For assays involving young adults, animals were age-matched based on the timing of completion of the L4 larval molt. All assays involving adult animals were performed using age-matched adult hermaphrodites 20-40 hours past the late L4 stage.

### Plasmid and strain construction

#### Calcium reporter transgenes

##### Vulval Muscle Ca^2+^

To visualize vulval muscle Ca^2+^ activity in adult animals, we used LX1918 *vsIs164 [unc-103e::GCaMP5::unc-54 3′UTR + unc-103e:: mCherry::unc-54 3′UTR + lin-15(+)] lite-1 (ce314) lin-15(n765ts) X* strain as described (Collins et al., 2016). In this strain, GCaMP5G (Akerboom et al., 2013) and mCherry are expressed from the *unc-103e* promoter (Collins and Koelle, 2013). The *unc-103e* promoter is only weakly expressed in vulval muscles during the L4 stages. To visualize vulval muscle activity in L4 animals, we expressed GCaMP5G and mCherry from the *ceh-24* promoter (Harfe and Fire, 1998). A ~2.8 kB DNA fragment upstream of the *ceh-24* start site was amplified from genomic DNA by PCR using the following oligonucleotides: 5’-GCG GCA TGC AAC GAG CCA TCC TAT ATC GGT GGT CCT CCG-3’ and 5’-CAT CCC GGG TTC CAA GGC AGA GAG CTG CTG-3’. This DNA fragment was ligated into pKMC257 (mCherry) and pKMC274 (GCaMP5G) from which the *unc-103e* promoter sequences were excised to generate pBR3 and pBR4, respectively. pBR3 (20 ng/μl) and pBR4 (80 ng/μl) were injected into LX1832 *lite-1(ce314) lin-15(n765ts) X* along with the pLI5EK rescue plasmid (50 ng/μl) (Clark et al., 1994). The extrachromosomal transgene produced was integrated by irradiation with UV light after treatment with trimethylpsoralen (UV/TMP) creating two independent transgenes *keyIs12* and *keyIs13*, which were then backcrossed to LX1832 parental line six times to generate the strains MIA51 and MIA53. Strain MIA51 *keyIs12 [ceh-24::GCaMP5::unc-54 3’UTR + ceh-24::mCherry::unc-54 3’UTR + lin-15(+)] IV; lite-1 (ce314) lin-15 (n765ts) X* was subsequently used for Ca^2+^ imaging. We noted repulsion between *keyIs12* and *wzIs30 IV*, a transgene that expresses Channelrhodopsin-2::YFP in HSN from the *egl-6* promoter (Emtage et al., 2012), suggesting both were linked to chromosome IV. As a result, we crossed MIA53 *keyIs13 [ceh-24::GCaMP5::unc-54 3’UTR + ceh-24::mCherry::unc-54 3’UTR + lin-15(+)]; lite-1 (ce314) lin-15(n765ts) X* with LX1836 *wzIs30 IV; lite-1(ce314) lin-15(n765ts) X*, generating MIA88 which was used to activate HSN neurons and record vulval muscle Ca^2+^ in L4 animals. In the case of young adults (3.5 & 6.5h post molt) and 24h old adults, strain LX1932 *wzIs30 IV; vsIs164 lite-1(ce314) lin-15(n765ts) X* was used as described (Collins et al., 2016).

##### HSN Ca^2+^

To visualize HSN Ca^2+^ activity in L4 and adult animals, we used the LX2004 *vsIs183* [*nlp-3::GCaMP5::nlp-3 3’UTR + nlp-3::mCherry::nlp-3 3’UTR + lin-15(+)] lite-1(ce314) lin-15(n765ts) X* strain expressing GCaMP5G and mCherry from the *nlp-3* promoter as previously described (Collins et al., 2016). In order to visualize HSN Ca^2+^ activity in *lin-12(wy750)* mutant animals lacking post-synaptic vm2 vulval muscle arms, we crossed MIA194 *lin-12(wy750) III* with LX2004 *vsIs183 lite-1(ce314) lin-15(n765ts) X* to generate MIA196 *lin-12(wy750)* III; *vsIs183 X lite-1(ce314) lin-15 (n765ts) X.* In order to visualize HSN Ca^2+^ activity in *glp-1(or178ts)* mutant animals, we crossed EU552 *glp-1(or178ts) III* with LX2004 *vsIs183 lite-1(ce314) lin-15(n765ts) X* to generate MIA219 *glp-1(or178ts) III; vsIs183 lite-1(ce314) lin-15(n765ts) X.*

#### Histamine gated chloride channel (HisCl) expressing transgenes

##### Vulval muscle HisCl

To produce a vulval muscle-specific HisCl transgene, coding sequences for mCherry in pBR3 were replaced with that for HisCl. First, an EagI restriction site (3’ of the mCherry encoding sequence) was changed to a NotI site using Quickchange site-directed mutagenesis to generate pBR5. The ~1.2 kB DNA fragment encoding the HisCl channel was amplified from pNP403 (Pokala et al., 2014) using the following oligonucleotides: 5’-GCG GCT AGC GTA GAA AAA ATG CAA AGC CCA ACT AGC AAA TTG G-3’ and 5’-GTG GCG GCC GCT TAT CAT AGG AAC GTT GTC-3’, cut with NheI/NotI, and ligated into pBR5 to generate pBR7. pBR7 (80 ng/μl) was injected into LX1832 along with pLI5EK (50 ng/μl). One line bearing an extrachromosomal transgene was integrated with UV/TMP, and six independent integrants *(keyIs14* to *keyIs19)* were recovered. Four of these were then backcrossed to the LX1832 parental line six times to generate strains MIA68, MIA69, MIA70, and MIA71. All four strains were used for behavioral assays in adult animals to test the effect of vulval muscle silencing on egg laying (Fig. 4B). MIA71 *keyIs19 [ceh-24::HisCl::unc-54 3’UTR + lin-15(+)]; lite-1(ce314) lin-15(n765ts) X* strain was used to study the effect of acute silencing of early activity on egg-laying behavior (Fig. 4C). To visualize HSN Ca^2+^ activity after vulval muscle silencing, we crossed MIA71 with LX2004 to generate strain MIA80 *keyIs19; vsIs183 lite-1(ce314) lin-15(n765ts) X.*

##### HSN HisCI

The ~1.2 kB DNA fragment encoding the HisCl channel was amplified from pNP403 using the following oligonucleotides: 5’-GCG GCT AGC GTA GAA AAA ATG CAA AGC CCA ACT AGC AAA TTG G-3’ and 5’-GCG GAG CTC TTA TCA TAG GAA CGT TGT CCA ATA GAC AAT A-3’. The amplicon was digested with NheI/SacI and ligated into similarly cut pSF169 (pegl-6::mCre (Flavell et al., 2013)) to generate pBR10. To follow expression in HSN, mCherry was amplified using the following oligonucleotides: 5’-GCG GCT AGC GTA GAA AAA ATG GTC TCA AAG GGT-3’ and 5’-GCG GAG CTC TCA GAT TTA CTT ATA CAA TTC ATC CAT G-3’. This amplicon was digested with NheI/SacI and ligated into pSF169 to generate pBR12. pBR10 (HisCl; 5 ng/μl) and pBR12 (mCherry; 10 ng/μl) were injected into LX1832 *lite-1 (ce314) lin-15(n765ts)* along with pLI5EK (50 ng/μl). The extrachromosomal transgene produced was integrated with UV/TMP, creating three independent integrants *(keyIs20, keyIs21, and keyIs22).* The resulting animals were backcrossed to the LX1832 parental line six times to generate strains MIA115, MIA116, and MIA117. The MIA116 strain had a low incidence of HSN developmental defects and was used subsequently for behavioral assays.

##### AII neuron HisCI

pNP403 was injected into LX1832 *lite-1(ce314) lin-15(n765ts)* animals at 50 ng/μl along with pLI5EK (50 ng/μl) to produce strain MIA60 carrying extrachromosomal transgene *keyEx16 [tag-168::HisCl::SL2::GFP + lin15(+)].* Non-Muv, *lin-15(+)* animals with strong GFP expression in the HSNs and other neurons were selected prior to behavioral silencing assays. All animals showed histamine-dependent paralysis that recovered after washout.

#### Transgenic reporters of circuit development and morphology

##### Vulval muscle morphology

To visualize vulval muscle development at the L4 stages, we injected pBR3 *[pceh-24::*mCherry] (80 ng/μl) along with a co-injection marker pCFJ90 (10 ng/μl) into TV201 *wyIs22* [punc-86::GFP::RAB-3 + podr-2::dsRed] (Patel et al., 2006) to generate an extrachromosomal transgene, *keyEx42.* To visualize adult vulval muscle morphology, we used the LX1918 *vsIs164 [unc-103e::GCaMP5::unc-54 3’UTR + unc-103e::mCherry::unc-54 3’UTR + lin-15(+)] lite-1 (ce314) lin-15(n765ts) X* strain (Collins et al., 2016). To visualize the expression of the *ser-4* gene, we used the strain AQ570 *[ijIs570]* (Tsalik and Hobert, 2003; Gurel et al., 2012).

##### HSN morphology

We used the LX2004 strain expressing mCherry from the *nlp-3* promoter to visualize HSN morphology at L4 stages as well as in adults. To visualize GFP::RAB-3 synaptic localization in HSNs during development, the *wyIs22* transgene was used (Patel et al., 2006).

##### Whole circuit morphology (HSN, VC and uv1 cells)

A ~3.2 kB DNA fragment upstream of the *ida-1* start site (Cai et al., 2004) was cloned using the following oligonucleotides: 5’-GCG GCA TGC CCT GCC TGT GCC AAC TTA CCT-3’ and 5’-CAT CCC GGG GCG GAT GAC ACA GAG ATG CGG-3’. The DNA fragment was digested with SphI/XmaI and ligated into pKMC257 and pKMC274 to generate plasmids pBR1 and pBR2. pBR1 (20 ng/μl) and pBR2 (80 ng/μl) were co-injected into LX1832 along with pLI5EK (50 ng/μl). The extrachromosomal transgene produced was integrated with UV/TMP creating four independent integrants *keyIs8* to *keyIs11*, which were then backcrossed to LX1832 parental line six times. MIA49 *keyIs11 [ida-1::GCaMP5::unc-54 3’UTR + ida-1::mCherry::unc-54 3’UTR + lin-15(+)]; lite-1(ce314) lin-15 (n765ts) X* was used subsequently to visualize whole-circuit morphology.

### Fluorescence imaging

#### 3D confocal microscopy

To visualize the morphological development of the egg-laying system, L4s and age-matched adults were immobilized using 10 mM muscimol on 4% agarose pads and covered with #1 coverslips. Two-channel confocal Z-stacks (along with a bright-field channel) using a pinhole opening of 1 Airy Unit (0.921 μm thick optical sections, 16-bit images) were obtained with an inverted Leica TCS SP5 confocal microscope with a 63X Water Apochromat objective (1.2NA). GFP and mCherry fluorescence was excited using a 488 nm and 561 nm laser lines, respectively. Images were analyzed in Volocity 6.3.1 (Perkin Elmer) and FIJI (Schindelin et al., 2012).

#### Ratiometric Ca^2+^ Imaging

Ratiometric Ca^2+^ recordings were performed on freely behaving animals mounted between a glass coverslip and chunk of NGM agar as previously described (Collins and Koelle, 2013; Li et al., 2013; Collins et al., 2016; Ravi et al., 2018). Recordings were collected on an inverted Leica TCS SP5 confocal microscope using the 8 kHz resonant scanner at ~20 fps at 256×256 pixel resolution, 12-bit depth and ≥2X digital zoom using a 20x Apochromat objective (0.7 NA) with the pinhole opened to ~20 μm. GCaMP5G and mCherry fluorescence was excited using a 488 nm and 561 nm laser lines, respectively. L4 animals at the relevant stages of vulval development were identified based on vulval morphology (Mok et al., 2015). Adult recordings were performed 24 hours after the late L4 stage. Young adults (3.5–6.5 h) were staged after cuticle shedding at the L4 to adult molt. After staging, animals were allowed to adapt for ~30 min before imaging. During imaging, the stage and focus were adjusted manually to keep the relevant cell/pre-synapse in view and in focus.

Ratiometric analysis (GCaMP5:mCherry) for all Ca^2+^ recordings was performed after background subtraction using Volocity 6.3.1 as described (Collins et al., 2016; Ravi et al., 2018). The egg-laying active state was operationally defined as the period one minute prior to the first egg-laying event and ending one minute after the last (in the case of a typical active phase where 3-4 eggs are laid in quick succession). However, in cases where two egg-laying events were apart by >60 s, peaks were considered to be in separate active phases and transients between these were considered to be from the inactive state.

#### Ratiometric Ca^2+^ comparisons with different reporters and developmental stages

To facilitate comparisons of ΔR/R between different reporters at different developmental stages, particularly during periods of elevated Ca^2+^ activity, HSN recordings in which baseline GCaMP5/mCherry fluorescence ratio values that were between 0.2-0.3 were selected for the analysis, while vulval muscle recordings with GCaMP5/mCherry ratio values between 0.1-0.2 were chosen (≥80% of recordings). Because HSN Ca^2+^ transient amplitude did not change significantly across developmental stages or in mutant or drug-treatment backgrounds, our analyses focused on HSN Ca^2+^ transient frequency. To test whether vulval muscle Ca^2+^ transient amplitudes recorded using different transgenes were suitable for quantitative comparisons, we measured the average GCaMP5:mCherry fluorescence ratio from two 15 by 15 μm regions of interest (ROI) from the anterior and posterior vulval muscles under identical imaging conditions (data not shown). The ROIs were positioned so as to ensure maximal coverage of the muscle cell area. We found that resting GCaMP5:mCherry ratios (±95% confidence intervals) bearing either the *ceh-24 (keyIs12)* or *unc-103e (vsIs164)* vulval muscle Ca^2+^ reporter transgenes were not statistically different at the developmental stages under comparison in Fig. 3H (L4.7-8 *(ceh-24):* 1.055 ±0.027; L4.9 *(ceh-24):* 1.055 ±0.061; Adult *(unc-103e):* 1.15 ±0.064; n≥10 animals measured per developmental stage). The coordination of vulval muscle contraction was determined as described (Li et al., 2013).

#### ERG expression analysis

To measure ERG *(unc-103e)* expression in the vulval muscles during development in staged LX1918 L4.7-8 and L4.9 larvae and 24-hour adults, we used identical imaging conditions to measure mCherry fluorescence through a 20x Plan Apochromat objective (0.8NA) using a Zeiss Axio Observer microscope onto a Hamamatsu ORCA Flash 4.0 V2 sCMOS sensor after excitation with a 590 nm LED (Zeiss Colibri.2). After import into Volocity, two 15 x 15 μm ROIs were placed on the anterior and posterior vulval muscles, and the mCherry fluorescence of the two objects was averaged. A control ROI placed outside of the animal was used for background subtraction.

### Behavior Assays and Microscopy

#### Optogenetics and Defecation Behavior Assays

ChR2 expressing strains were maintained on OP50 with or without *all-trans* retinal (ATR) (0.4 mM). ChR2 was activated during Ca^2+^ imaging experiments with the same, continuous laser light used to excite GCaMP5 fluorescence.

#### Acute silencing experiments using HisCl

For acute silencing assays, NGM plates containing 10 mM histamine were prepared and used as described (Pokala et al., 2014). For adult behavioral assays, HisCl expressing strains were staged as late L4s with histamine treatment and behavior assays performed 24 hours later. For L4 activity silencing, L4.7 animals were placed on NGM plates with or without 10 mM histamine and were monitored to note when the animals complete the L4 molt. Each animal was then transferred to a new seeded plate (lacking histamine), and the time for each animal to lay its first egg was recorded.

#### Animal sterilization

Animals were sterilized using Floxuridine (FUDR) as follows. 100 μl of 10 mg/ml FUDR was applied to OP50 seeded NGM plates. Late L4 animals were then staged onto the FUDR plates and the treated adults were imaged 24 hours later. MIA219 *glp-1(or178ts) III; vsIs183 lite-1(ce314) lin-15(n765ts) X* animals were sterilized during embryogenesis as described (Fujiwara et al., 2016). L1-L2 animals were shifted to 25 °C and returned to 15 °C after 24 hours. Late L4 animals were then staged and grown at 15°C and imaged 24 hours later.

### Experimental Design and Statistical Analysis

Sample sizes for behavioral assays followed previous studies (Chase et al., 2004; Collins and Koelle, 2013; Collins et al., 2016). No explicit power analysis was performed before the study. Statistical analysis was performed using Prism 6 (GraphPad). Ca^2+^ transient peak amplitudes, widths, and inter-transient intervals were pooled from multiple animals (typically ~10 animals per genotype/condition per experiment). No animals or data were excluded except as indicated above to facilitate comparisons of Ca^2+^ transient amplitudes between different development stages and reporters. Individual *p* values are indicated in each Figure legend, and all tests were corrected for multiple comparisons (Bonferroni for ANOVA; Dunn for Kruskal-Wallis).

## Results

### Asynchronous presynaptic and postsynaptic development in the *C. elegans* egg-laying behavior circuit

We have previously described the function of cell activity in the adult egg-laying behavior circuit and how developmental mutations impact circuit activity and adult behavior (Collins and Koelle, 2013; Li et al., 2013; Collins et al., 2016). Because development of the cells in the circuit is known to be complete by the end of the fourth larval (L4) stage (Li and Chalfie, 1990), we wanted to determine the relationship between circuit development and the emergence of cell activity as the animals mature from juveniles into egg-laying adults. We exploited the stereotyped morphology of the developing primary and secondary vulval epithelial cells in the fourth (final) larval stage to define discrete half-hour stages of development until the L4-adult molt (Fig. 1A-F) as described (Mok et al., 2015). We observed NLP-3 neuropeptide promoter expression in HSNs of late L4 animals (Fig. G-I), showing that L4.7-8 HSNs have specified a transmitter phenotype. Consistent with L4.7-8 HSN being functional, the presynaptic marker GFP::RAB-3 expressed from the *unc-86* promoter showed clear punctate localization in HSN at synaptic sites at these stages (Fig. 1J-L), confirming previous observations with light microscopy and serial electron microscopy reconstruction that HSN development is complete by L4.7-8 (Shen and Bargmann, 2003; Shen et al., 2004; Adler et al., 2006; Patel et al., 2006).

**Fig. 1:**
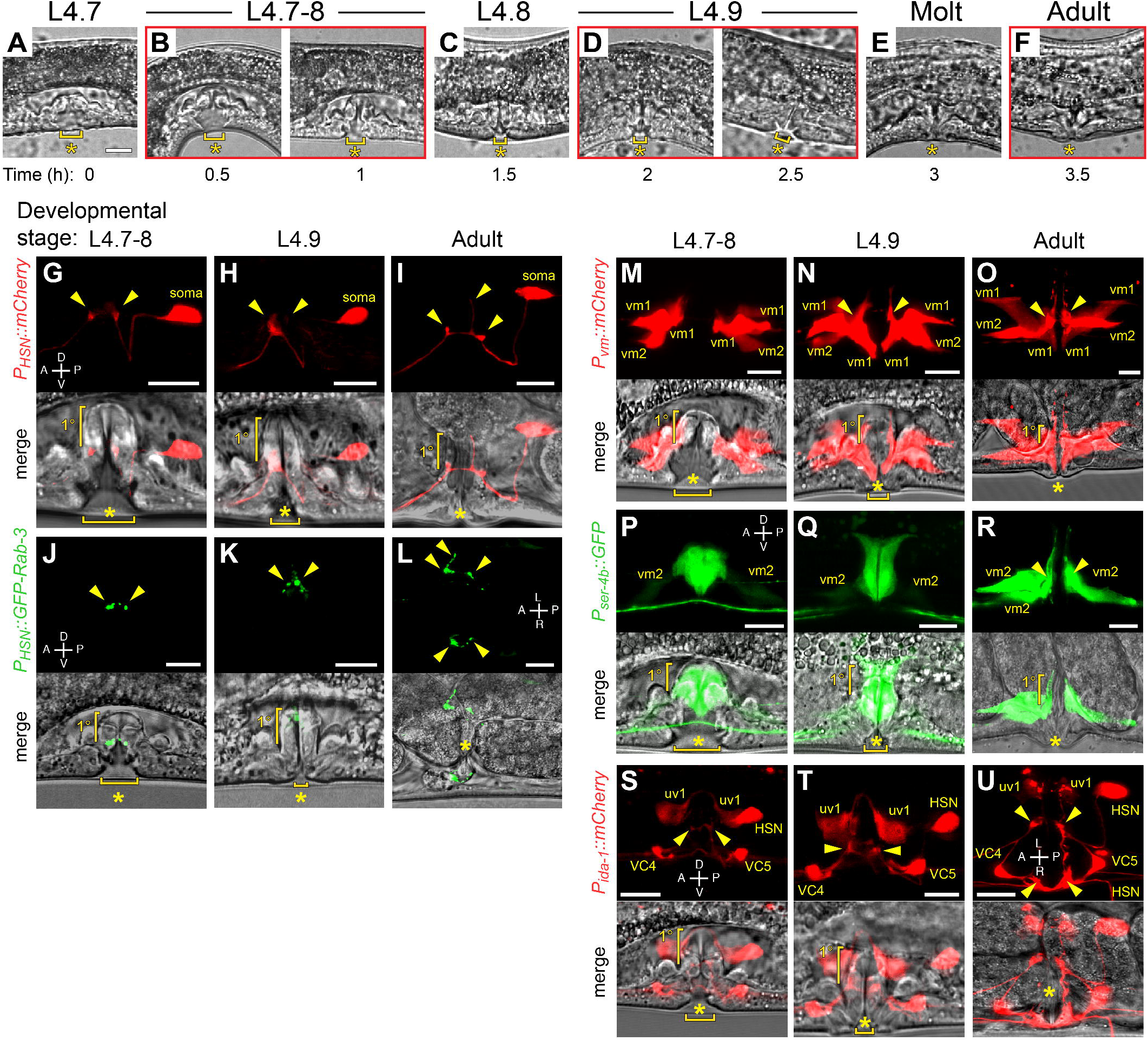
Morphological development of the *C. elegans* egg-laying circuit. (A-F) Representative images of vulval morphology at late L4 stages-(A) L4.7, (B) L4.7-8, (C) L4.8, (D) L4.9, (E) Molt and (F) Young adult. (G-I) Morphology of HSN labeled with mCherry (top) and the vulva (bottom) in L4.7-8 (G) and L4.9 (H) larval stages and in adults (I). (J-L) Morphology of HSN synapses labeled with GFP::RAB-3 (top) and the vulva (bottom) in L4.7-8 (J) and L4.9 (K) larval stages and in adults (L). (M-O) Morphology of vm1 and vm2 vulval muscles labeled with mCherry (top) and the vulva (bottom) in L4.7-8 (M) and L4.9 (N) larval stages and in adults (O). (P-R) Developmental expression of *ser-4* from a GFP transcriptional reporter (top) at the L4.7-8 (P) and L4.9 (Q) larval stages and in adults (R) and the vulva (bottom). (S-U) Morphology of HSN, VC4, VC5, and the uv1 neuroendocrine cells labeled with mCherry (top) and the vulva (bottom) in L4.7-8 (S) and L4.9 (T) larval stages and in adults (U) visualized using the *ida-1* promoter. Arrowheads in all images indicate the location of presynaptic boutons or postsynaptic vm2 muscle arms. Scale bar is 10 μm; anterior is at left and ventral is at bottom unless indicated otherwise. Asterisks indicate the position of the developing or completed vulval opening. Vertical half-brackets indicate the approximate position of primary (1°) vulval epithelial cells, and horizontal bracket indicates progress of vulval lumen collapse at each larval stage.

Unlike HSNs, we found the post-synaptic vulval muscles completed their morphological development during the L4.9 stage, just prior to the L4 molt. We expressed mCherry in the vulval muscles from the *ceh-24* promoter (Harfe and Fire, 1998) and found that the vm1 and vm2 vulval muscles were still developing at the L4.7-8 stage (Fig. 1M). After lumen collapse at the L4.9 stage, the tips of the vm1 muscles extended ventrally to the lips of the vulva, and the anterior and posterior vm2 muscle arms extended laterally along the junction between the primary and secondary vulval epithelial cells (Fig. 1N), making contact with each other at the HSN (and VC) synaptic release sites that continues in adults (Fig. 10). Previous work has shown that mutations that disrupt LIN-12/Notch signaling perturb development of the vm2 muscle arms in late L4 animals (Li et al., 2013), a time when we observed vm2 muscle arm extension.

Vulval muscles express multiple serotonin receptors that mediate their response to HSN input (Carnell et al., 2005; Dempsey et al., 2005; Hobson et al., 2006; Hapiak et al., 2009) In order to look at the developmental expression pattern of one such serotonin receptor, we examined a transgenic reporter line expressing GFP under the *ser-4b* gene promoter (Tsalik and Hobert, 2003; Gurel et al., 2012). As shown in Fig. 1P and 1Q, we observed strong GFP expression in VulF and VulE primary and VulD secondary epithelial cells. The *ser-4b* promoter also drove weak GFP expression in the vm2 muscles in L4.7-9, and this was elevated in adults (Fig. 1P-R). Serial EM reconstruction has shown that HSN makes transient synapses onto the vulval epithelial cells in developing L4 animals, and the expression of a serotonin receptor in these cells and the vm2 muscles during this period suggests they have specified a receptor phenotype (Shen et al., 2004). Lastly, we wanted to determine whether the VC motor neurons and uv1 neuroendocrine cells had completed their development in late L4 animals. To simultaneously visualize HSN, VC, and the uv1 neuroendocrine cells, we expressed mCherry from the *ida-1* promoter, a gene expressed in a subset of peptidergic cells, including those in the egg-laying circuit (Cai et al., 2004). We observed mCherrry expression in all three cell types in L4.7-8 animals, consistent with their development of a peptidergic phenotype in late L4 animals (Fig. 1S-U). As expected, HSN and VC presynaptic termini assembled at the junction between the primary and secondary vulval epithelial cells in L4.7-8. The uv1 cells were positioned laterally to the HSN/VC synaptic regions and extended dorsal processes around the primary vulval epithelial cells (Fig.1S-U). These results indicate that the morphological development and peptidergic expression phenotype of the HSN, VC, and uv1 cells is largely complete by L4.7-8 stage. In contrast, vulval muscle morphological development is completed in the L4.9 stage when the vm2 muscle arms reach each other and the HSN and the VC presynaptic boutons and begin to express the serotonin receptor SER-4b.

### HSNs switch from tonic activity in juveniles to burst firing in egg-laying adults

We next wanted to determine if the HSNs show activity as they develop and how that activity compares to that seen in egg-laying adults. To follow HSN activity, we expressed the Ca^2+^ reporter GCaMP5 along with mCherry in HSN using the *nlp-3* promoter and performed ratiometric Ca^2+^ imaging in freely behaving animals as previously described (Collins et al., 2016). Starting at the L4.7-8 larval stage, we observed rhythmic Ca^2+^ activity in both HSN presynaptic termini and in the soma (Fig. 2A and 2B). During the L4.9 larval stage, when animals exhibited behavioral features of the developmentally timed L4 quiescence (Raizen et al., 2008), rhythmic Ca^2+^ activity in the HSNs slowed (Fig. 2B; Movie 1). The tonic HSN activity we observed in juveniles (Fig. 2B; Movie 2) differed from the alternating, two-state pattern previously seen in adult animals where periods of infrequent activity are interrupted by bouts of HSN burst firing that drive the egg-laying active state (Collins et al., 2016). We quantitated changes in HSN Ca^2+^ transient peak amplitude and frequency during the different developmental stages and behavior states. We found no significant differences in HSN Ca^2+^ transient amplitude (Fig. 2C), but we did observe significant changes in frequency. The median inter-transient interval in L4.7-8 animals was ~34 s, and this interval increased to ~60 s as animals reached the L4.9 stage (Fig. 2D). The reduction of HSN transient frequency seen in L4.9 animals resembled the egg-laying inactive state. However, none of the developmental stages recapitulated the ‘burst’ Ca^2+^ activity with <20 s inter-transient intervals seen during the egg-laying active state (Fig. 2D). Together, these results indicate that the HSNs show tonic Ca^2+^ activity after their morphological development is complete. HSN activity then switches into distinct inactive and active states as animals become egg-laying adults.

**Fig. 2.**
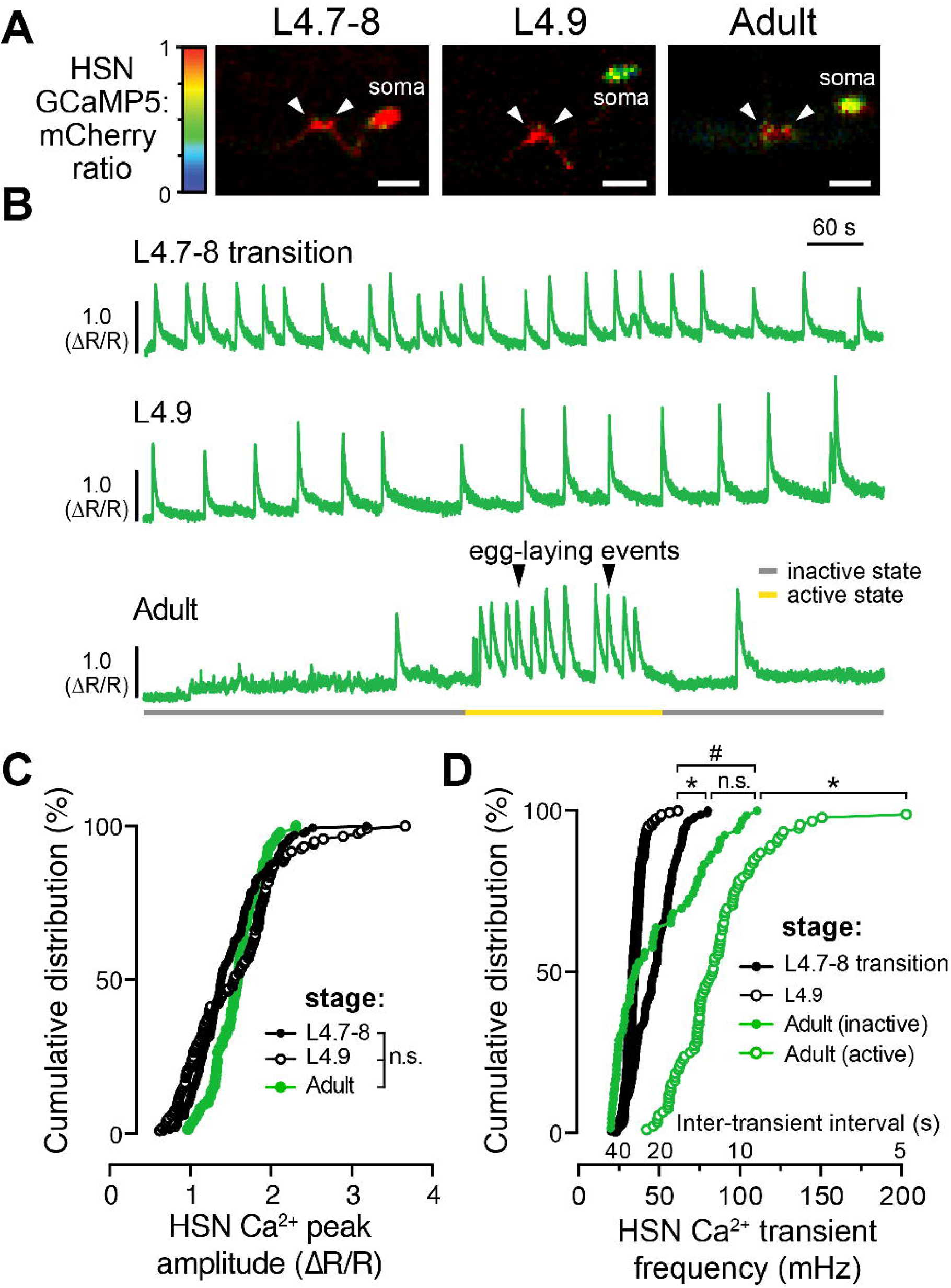
HSN neurons show tonic Ca^2+^ activity during the late L4 stage and burst firing during the egg-laying active state. (A) Representative images of the intensity-modulated GCaMP5:mCherry fluorescence ratio during HSN Ca^2+^ transients in L4.7-8 and L4.9 larval stages, and in adults. White arrowheads show Ca^2+^ activity localized to the anterior and posterior presynaptic boutons. Scale bar is 10 μm; anterior is at left, ventral is at bottom. See also Movies 1 and 2. (B) Representative GCaMP5: mCherry ratio traces (ΔR/R) of HSN Ca^2+^ activity in L4.7-8 (top), L4.9 (middle), and in adult animals (bottom). Adults show distinct active (yellow) and inactive (grey) egg-laying behavior states. Black arrowheads indicate egg-laying events. (C) Cumulative distributions of HSN Ca^2+^ peak amplitudes in L4.7-8 (filled black circles), L4.9 (open black circles), and adults (filled green circles). n.s. indicates p>0.0809 (one-way ANOVA). (D) Cumulative distribution plots of instantaneous HSN Ca^2+^ transient frequencies (and inter-transient intervals) from L4.7-8 (filled black circles) and L4.9 (open black circles) animals, and from adult egg-laying inactive (filled green circles) and active (green open circles) states. Asterisks (*) indicate p<0.0001; pound sign (#) indicates p=0.0283; n.s. indicates p=0.1831 (Kruskal-Wallis test).

The onset of Ca^2+^ activity in the HSN neurons during the late L4 stage coincided with changes in animal locomotion, pharyngeal pumping, and defecation behaviors that accompany the L4 lethargus (Raizen et al., 2008). Previous published work has shown that there is an increase in animal locomotion in adult animals around egg-laying active states driven by serotonin signaling from HSN onto AVF (Hardaker et al., 2001). Loss of HSN neurons or serotonin signaling from HSN reduces reversals and increases forward locomotion and exploratory behavior (Flavell et al., 2013). To understand if the tonic HSN activity seen in juveniles was associated with locomotor arousal, we analyzed movement in L4.9 animals ten seconds before and after each HSN Ca^2+^ transient. About one third of L4.9 HSN transients failed to show any movement before or after the transient (35±7%), and the remaining HSN transients were about evenly split between those which showed movements before (30±7%), after (15±7%), or before and after (20±7%) the transient (n=156 transients). These results show that although HSN Ca^2+^ transients can occur around locomotion events, there does not appear to be a causal relationship between HSN activity and movement in juvenile animals.

### Vulval muscle Ca^2+^ transients increase in strength and frequency during development

We next wanted to determine if the HSN activity we observe in late L4 animals drives early vulval muscle activity. We used the *ceh-24* promoter to drive expression of GCaMP5 and mCherry in the vulval muscles of L4 animals. We detected Ca^2+^ transients at the L4.7-8 larval stage in the still-developing vulval muscles, and these transients continued and increased in frequency as the muscles completed their development at the L4.9 stage (Fig. 3A-C, 3F and 3G; Movies 3-5). The median interval between vulval muscle Ca^2+^ transients was ~32 s in L4.7-8 animals which dropped to 18 s in L4.9 animals. L4 vulval muscle activity differs from that observed previously in egg-laying adults (Fig. 3D and 3E; Movie 6). The frequency of vulval muscle Ca^2+^ transients increased significantly in animals during the egg-laying active state with median intervals dropping to ~7 s phased with each body bend (Fig. 3G), as previously described (Collins and Koelle, 2013; Collins et al., 2016). We found that vulval muscle Ca^2+^ transients become stronger after development. While Ca^2+^ transient amplitudes in the L4.7-8 and L4.9 stages were not significantly different, inactive phase Ca^2+^ transients of adults were stronger than those observed in L4 animals (Fig. 3H). In adult animals, strong Ca^2+^ transients were observed during the egg-laying active states, with the strongest Ca^2+^ transients driving the complete and simultaneous contraction of anterior and posterior vulval muscles to allow egg release (Fig. 3E and 3H).

**Fig. 3.**
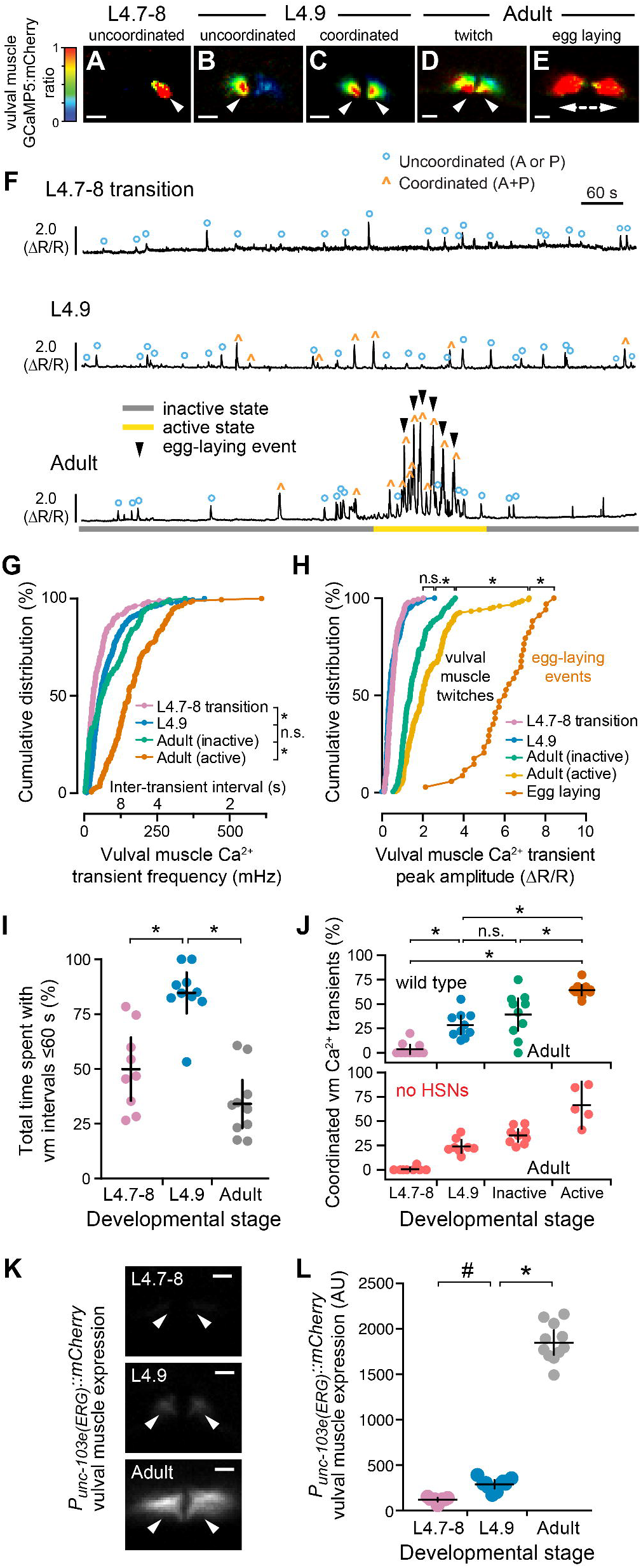
Development of coordinated vulval muscle Ca^2+^ transients in the L4.9 stage does not require presynaptic HSN input. (A-E) Representative images of the intensity-modulated GCaMP5:mCherry fluorescence ratio during vulval muscle Ca^2+^ transients at the L4.7-8 (A), L4.9 larval stages (B,C), and during the adult active state (D,E). White arrowheads show localization of Ca^2+^ transients. Scale bars are 10 μm; anterior at left, ventral at bottom. See also Movies 3-6. (F) Representative GCaMP5:mCherry (ΔR/R) ratio traces of vulval muscle Ca^2+^ activity at L4.7-8 (top), L4.9 (middle), and in adult animals (bottom) during an inactive (grey) and active (yellow) egg-laying state. Uncoordinated transients are indicated by blue circles (°), coordinated transients by orange carets (^), egg-laying events by black arrowheads. (G and H) Cumulative distribution plots of instantaneous vulval muscle Ca^2+^ transient peak frequencies (G) and amplitudes (H) at L4.7-8 (pink), L4.9 (blue), and in the egg-laying inactive (green) and active state (orange) of adults. Asterisks indicate *p*<0.0001; n.s. indicates *p*>0.9999 (Kruskal-Wallis test). (I) Scatterplots show time spent by 9-10 animals with frequent Ca^2+^ transients (inter-transient intervals ≤60 s) at L4.7-8 (pink), L4.9 (blue), and in adults (gray). Error bars show 95% confidence interval for the mean. Asterisks indicates *p*≤0.0002 (one-way ANOVA). (J) Scatterplots show percent synchronous anterior and posterior vulval muscle Ca^2+^ transients in each individual at L4.7-8 (pink), L4.9 (blue), and in adult egg-laying inactive (green) and active states (orange) in wildtype (top) and *egl-1(n986dm)* animals (red) lacking HSNs (bottom). Error bars show 95% confidence intervals for the mean from ≥5 animals. Asterisks indicate *p*≤0.0022; n.s. indicates p≥0.1653 (one-way ANOVA). (K) Representative images of mCherry fluorescence in the vulval muscles from a *unc-103e* (ERG) transcriptional reporter in an L4.7-8, L4.9, and adult animal. White arrowheads show anterior (left) and posterior (right) vulval muscle cells; scale bar is 10 μm. (L) Scatterplots show mCherry fluorescence from the *unc-103e* promoter in ten animals. Error bars show 95% confidence interval for the mean; ‘#’ indicates *p*=0.0288 and asterisk indicates *p*≤0.0001 (one-way ANOVA).

We were surprised that vulval muscle transient frequencies decreased in adults as circuit activity bifurcated into distinct inactive and active egg-laying behavior states. We quantified periods of increased activity by measuring time spent with vulval muscle Ca^2+^ transient intervals less than one minute. We found that vulval muscle activity increased as L4.7-8 animals developed into L4.9 animals but then dropped significantly in egg-laying adults. L4.7-8 animals on average spent ~50% of their time in periods of increased vulval muscle activity, and this increased to 85% as animals entered the L4.9 stage (Fig. 3I). In contrast, adult animals spent only about ~33% of their time in periods with elevated vulval muscle activity (Fig. 3I) about half of which were coincident with the ~3 minute egg-laying active states that occur about every 20 minutes (Waggoner et al., 1998). What depresses vulval muscle activity in adult animals? We have previously shown that the loss of *unc-103*, which encodes Ether-a-Go-Go Related Gene (ERG) K+ channel, results in increased vulval muscle excitability and egg-laying behavior (Collins and Koelle, 2013). Using an mCherry transcriptional reporter transgene, we found that *unc-103e* expression in vulval muscles is low in L4 animals and increases >15-fold as animals mature into egg-laying adults (Fig. 3K and 3L). These results are consistent with our previous functional results that show that ERG depresses vulval muscle electrical excitability in adults to promote distinct inactive and active egg-laying behavior states (Collins and Koelle, 2013).

### Development of coordinated vulval muscle activity for egg laying

Egg release through the vulva requires the synchronous contraction of the anterior (A) and posterior (P) vulval muscles (Fig. 3E). Previous work has shown that loss of Notch signaling blocks postsynaptic vm2 muscle arm development in L4 animals resulting in asynchronous vulval muscle contractility and defects in egg-release in adults (Li et al., 2013). Because of the vulval slit, the lateral vm2 muscle arms that develop between L4.7-8 and L4.9 form the only sites of potential contact between the anterior and posterior vulval muscles (Fig. 1M and 1N). To determine the relationship between vulval muscle morphology and activity, we examined the spatial distribution of vulval muscle Ca^2+^ during identified transients. We found that only 5% of vulval muscle Ca^2+^ transients were coordinated in the L4.7-8 stage (Fig. 3A; Movie 3), with nearly all transients occurring in either the anterior or posterior muscles (Fig. 3F and 3J). The degree of vulval muscle coordination increased significantly to ~28% of transients during L4.9 (Fig. 3J; compare Movies 4 and 5) a time when vm1 and vm2 muscles, as well as vm2 muscle arms, complete their development (compare Fig. 1M and 1N). This level of coordinated muscle activity was not significantly different to that found in adult animals during the egg-laying inactive state (Fig. 3J; compare Fig. 3C and 3D). During the egg-laying active state ~60% of vulval muscle transients were found to be coordinated, with Ca^2+^ transients occurring synchronously in the anterior and posterior muscles (Movie 6).

To test whether HSN activity was required for the development of coordinated muscle activity, we analyzed muscle activity in animals missing the HSNs. Surprisingly, we observed that vulval muscles develop wild-type levels of coordinated activity even without HSN input (Fig. 3J). We have previously shown that vulval muscle activity in adults is phased with locomotion (Collins et al., 2016), possibly via rhythmic acetylcholine release from the VA7 and VB6 motor neurons that synapse onto the vm1 muscles (White et al., 1986). Vulval muscle activity in L4.9 animals accompanied ongoing locomotion as well. We analyzed recordings from L4.9 animals for movement ten seconds before and after each vulval muscle Ca^2+^ transient. A clear majority of transients (62±5%) were accompanied by movements occurring both before and after vulval muscle activity, with a smaller fraction of transients occurring just before or just after movement (11±4% and 10±4%, respectively; n=291 transients). Movement was not strictly required for vulval muscle activity, as Ca^2+^ transients were still observed in non-moving animals (17±4%). Our results show that coordinated vulval muscle activity in L4.9 stage is independent of HSN input and may instead be driven by input from the locomotion motor neurons into vm1 and through the lateral vm2 muscle contact along the vulval slit.

### Early neuronal and vulval muscle activity is not required for the onset of adult egg-laying behavior

Activity in developing circuits has previously been shown to contribute to mature patterns of activity that drive behavior. Is the early activity we observe in HSN and vulval muscles required for the proper onset of egg-laying behavior in adults? To test this, we first set out to determine when adults initiate egg laying. We found wild-type animals laid their first egg at about ~6-7 hours after the L4-adult molt (Fig. 4A) after accumulating ~8-10 eggs in the uterus, a time when VC and uv1 Ca^2+^ activity is first observed (data not shown). Animals without HSNs laid their first egg much later, ~18 hours post molt (Fig. 4A). Gain-of-function receptor mutations in EGL-6, a neuropeptide receptor coupled to Gα_O_ (Ringstad and Horvitz, 2008), or EGL-47, a putative gustatory receptor block neurotransmitter release from HSN (Moresco and Koelle, 2004) and delay egg release until ~15-17 hours after the L4 molt, resembling animals without HSNs (Fig. 4A). Surprisingly, tryptophan hydroxylase *(tph-1)* knockout animals that are unable to synthesize serotonin showed only a small albeit significant delay in egg release compared to wild type (~7-8 hours post L4 molt), suggesting that HSN promotes egg laying via release of neurotransmitters other than serotonin.

**Fig. 4:**
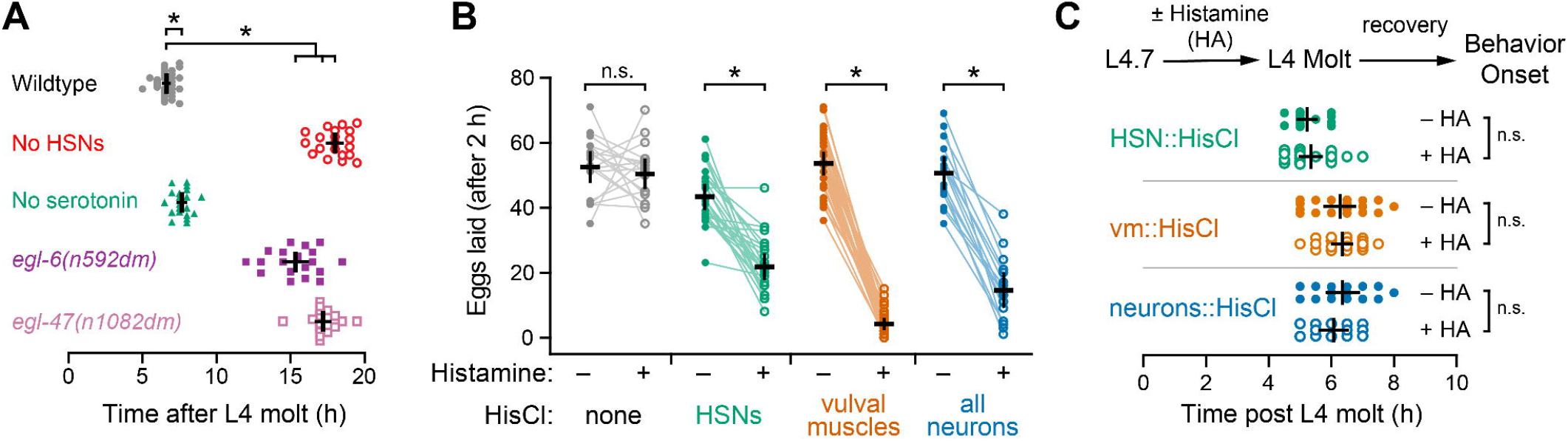
Early HSN and vulval muscle activity is not required for the onset of egg-laying behavior. (A) Scatter plots of the first egg-laying event in wild-type (grey), HSN-deficient *egl-1(n986dm)* (red open circles), serotonin-deficient *tph-1(mg280)* (green triangles), *egl-6(n592dm)* (purple squares), and *egl-47(n1082dm)* (pink open squares) mutant animals. Error bars show 95% confidence intervals for the mean from ≥19 animals. Asterisks indicate *p*≤0.0016 (One-way ANOVA). (B) Scatter plots showing eggs laid by three 24-hour adult animals in two hours before (filled circles) and in two hours after incubation on plates with 10 mM histamine (open circles). Transgenic animals expressing HisCl in vulval muscles (orange), HSN neurons (green), all neurons (blue) were compared with the non-transgenic wild-type (grey). Error bars indicate 95% confidence intervals for the mean from ≥17 paired replicates. Asterisks indicate *p*<0.0001; n.s. indicate *p*=0.5224 (paired Student’s t test). (C) Top, transgenic L4.7 animals expressing HisCl channels were incubated on NGM plates with or without 10 mM histamine until the L4-Adult molt. Animals were then moved to plates lacking histamine and allowed to recover and lay eggs. Bottom, scatter plots show the timing of the first egg-laying event with (open circles) and without (filled circles) histamine. Error bars indicate 95% confidence intervals for the mean; n.s. indicates *p*>0.9999 (one-way ANOVA).

To silence HSN and vulval muscle activity acutely and reversibly, we expressed *Drosophila* Histamine-gated chloride channels (HisCl) using cell-specific promoters and tested how histamine affected egg-laying behavior (Pokala et al., 2014). Egg laying was unaffected by exogenous histamine in non-transgenic animals but was potently inhibited when HisCl channels were transgenically expressed in the HSNs, the vulval muscles, or in the entire nervous system (Fig. 4B). Silencing these cells in late L4 animals for the entire period where we observe early activity caused no significant changes in the onset of adult egg laying after histamine washout in molted adults (Fig. 4C). We also observed no change in the steady-state number of unlaid eggs in the uterus after developmental silencing of L4 animals with histamine (data not shown). These results suggest that presynaptic and postsynaptic activity in the developing circuit is not required for circuit development or behavior.

### Vulval muscle responsiveness to HSN activity increases as maturing animals accumulate unlaid eggs

We and others have previously shown that optogenetic activation of the HSNs in adult animals is sufficient to induce egg-laying circuit activity and behavior (Emtage et al., 2012; Collins et al., 2016). Despite the fact that both the HSNs and vulval muscles show activity in L4.9 animals, egg laying does not begin until 6-7 hours later when the animals have accumulated ~8-10 unlaid eggs in the uterus. In order to dissect the relationship between developmental time, egg production, and circuit functionality, we tested when the vulval muscles develop sensitivity to HSN input. We optogenetically activated the HSNs using Channelrhodopsin-2 (ChR2) while simultaneously recording Ca^2+^ activity in the vulval muscles at 3 stages: in L4.9 juveniles and in 3.5 hour and 6.5-hour old adults. L4.9 animals have no eggs in the uterus, 3.5-hour adults contained 0-1 unlaid eggs, while 6.5-hour old adults had accumulated ~8-10 eggs. Stimulating HSNs in L4.9 juveniles or in 3.5-hour adults failed to induce detectable changes in vulval muscle Ca^2+^ activity (Fig. 5A, 5B, 5F). In contrast, optogenetic activation of HSNs in 6.5-hour adults significantly increased vulval muscle Ca^2+^ activity and triggered egg laying (Fig. 5C and 5F). L4.9 juveniles or 3.5-hour adults with 0-1 eggs in the uterus had a mean transient frequency of ≤100 mHz, similar to the inactive state vulval muscle Ca^2+^ response seen in 6.5-hour adult animals with ~8 unlaid eggs grown in the absence of ATR, a cofactor necessary for ChR2 activation. The vulval muscle Ca^2+^ response to HSN input was increased to ~170 mHz in 6.5-hour adults that had accumulated ~8 unlaid eggs (Fig. 5G). Surprisingly, vulval muscles in serotonin-deficient mutants responded normally to HSN activation at 6.5 hours (Fig. 5D and 5E), a finding consistent with the normal onset of egg laying in these mutants (Fig. 4A). Together, these results show that despite having significant Ca^2+^ activity in juveniles, the adult vulval muscles only develop a robust response to HSN input ~6 hours after the molt, a time when fertilized embryos are being deposited in the uterus to be laid.

**Fig. 5.**
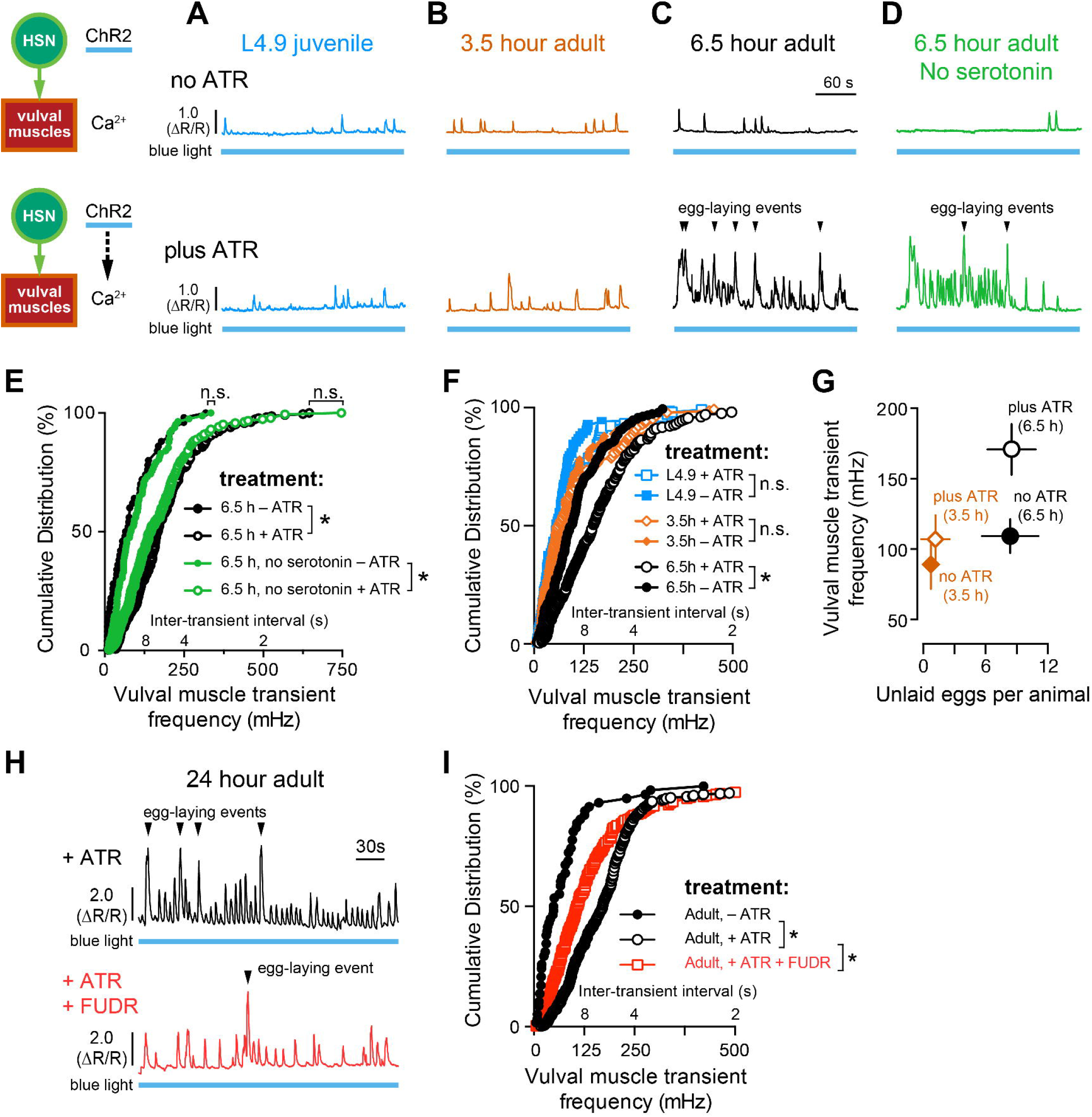
Vulval muscle responsiveness to HSN input correlates with egg accumulation. (A-D) Representative traces of vulval muscle Ca^2+^ activity in L4.9 juveniles (A, blue), 3.5-hour adults (B, orange), 6.5-hour wild-type adults (C, black), and 6.5-hour serotonin-deficient *tph-1(mg280)* mutant adults (D, green) with and without optogenetic activation of HSN. Animals were grown in the presence (plus ATR, top) or absence (no ATR, bottom) of *all-trans* retinal (see cartoon schematic). Continuous 489 nm laser light was used to simultaneously stimulate HSN ChR2 activity and excite GCaMP5 fluorescence for the entire recording. Arrowheads indicate egg-laying events. Blue bars under the Ca^2+^ traces indicate the period of continuous blue light exposure. (E) Cumulative distribution plots of instantaneous peak frequencies (and inter-transient intervals) of vulval muscle Ca^2+^ activity in 6.5-hour adult wild-type (black filled circles, no ATR; black open circles, plus ATR) and *tph-1(mg280)* mutant animals (green filled circles, no ATR; green open circles, plus ATR). Asterisks indicate *p*<0.0001; n.s. indicates p≥0.2863 (Kruskal-Wallis test). (F) Cumulative distribution plots of instantaneous peak frequencies (and inter-transient intervals) of vulval muscle Ca^2+^ activity in L4.9 juveniles (blue filled squares, no ATR; blue open squares, plus ATR), 3.5-hour old adults (orange filled circles, no ATR; orange open circles, plus ATR), and 6.5-hour old adults (black filled circles, no ATR; black open circles, plus ATR). Asterisk indicates *p*<0.0001; n.s. indicates p≥0.3836 (Kruskal-Wallis test). (G) Plot shows the average number of unlaid eggs present in the uterus and the average vulval muscle Ca^2+^ transient peak frequency, ±95% confidence intervals. (H) Representative traces of HSN-induced vulval muscle Ca^2+^ activity in untreated (top, black) and FUDR-treated 24-hour adult animals (bottom, red). Arrowheads indicate egg-laying events. (I) Cumulative distribution plots of instantaneous peak frequencies (and inter-transient intervals) of 2+ vulval muscle Ca^2+^ activity after optogenetic activation of HSNs in untreated animals grown with ATR (+ATR, open black circles), FUDR-treated animals with ATR (+ATR, open red circles), and in untreated animals without ATR (no ATR, filled black circles). Asterisks indicate *p*<0.0001 (Kruskal-Wallis test).

We next examined whether this change in vulval response in older adults was caused by ongoing developmental events or was instead a consequence of egg accumulation. We previously demonstrated that adults sterilized with FUDR, a chemical blocker of germline cell division and egg production, showed inactive state levels of vulval muscle activity (Collins et al., 2016). We found that vulval muscles in FUDR-treated animals 24 hours after the molt were also significantly less responsive to HSN optogenetic stimulation (Fig. 5H and 5I). The residual vulval muscle response in FUDR-treated animals is likely caused by incomplete sterilization when FUDR is added to L4.9 animals. We interpret these results as indicating that animal age or circuit maturity are not sufficient for the onset of the egg-laying active state.

### A retrograde signal of egg accumulation and vulval muscle activity drives presynaptic HSN activity

HSN activity can be inhibited by external sensory signals and feedback of egg release (Ringstad and Horvitz, 2008; Emtage et al., 2012; Collins et al., 2016; Banerjee et al., 2017), but the factors that promote HSN activity are not clear. We tested whether egg accumulation promotes circuit activity through the presynaptic HSNs, the postsynaptic vulval muscles, or both. We found that HSN Ca^2+^ activity, particularly the burst firing activity associated with the active state, was dramatically reduced in FUDR-treated animals (Fig. 6A). Although we did observe single HSN Ca^2+^ transients after FUDR treatment, the intervals in between were prolonged, often minutes apart (Fig. 6C). We quantified the total time spent by animals with HSN Ca^2+^ transient intervals <30 s apart as a measure of HSN burst-firing seen in the active state. We found that while untreated animals spent ~13% of their time with the HSNs showing high-frequency activity, such bursts were eliminated in FUDR-treated animals (Fig. 6D). We confirmed the FUDR results using a conditional *glp-1(or178ts)* Notch receptor mutant that causes germ line loss and sterility when shifted to 25°C during the L1 stage (Fig. 6B). We observed a dramatic reduction in HSN Ca^2+^ transient frequency in sterile *glp-1(or178ts)* adults, phenocopying the results seen with FUDR (Fig. 6B and 6C). While *glp-1(or178ts)* fertile animals (raised at 15°C) animals spent a typical 13% of their time with the HSNs showing high frequency activity, such bursts were eliminated in sterile *glp-1(or178ts)* adults (Fig. 6D). These results show that feedback of germline activity, egg production, and/or egg accumulation modulates the frequency of HSN activity.

**Fig. 6:**
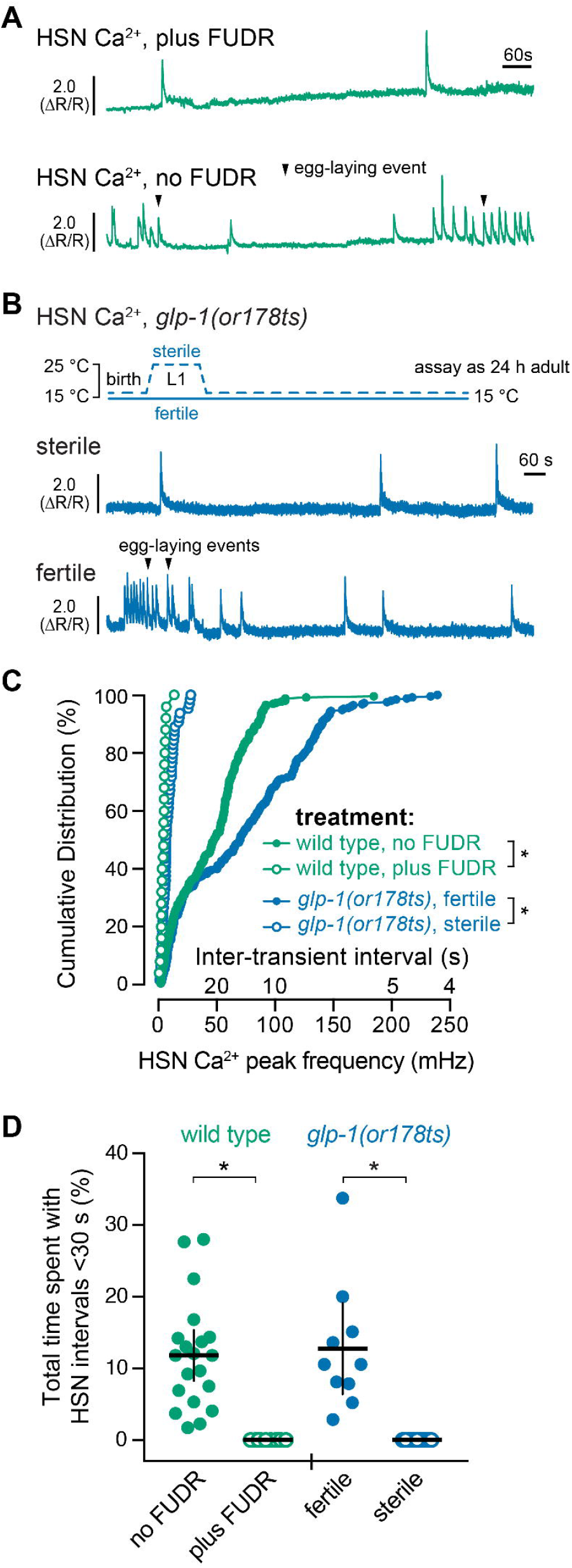
Germ line activity is required for HSN burst firing and the active state. (A) Representative HSN Ca^2+^ traces in untreated (top) and FUDR-treated (bottom) adult animals. (B) Representative HSN Ca^2+^ traces in adult *glp-1(or178ts)* sterilized animals (L1s shifted to 25°C for 24h and raised to adults at 15°C) (top) and *glp-1(or178ts)* fertile animals (raised at 15°C) (bottom). Arrowheads indicate egg laying events. (C) Cumulative distribution plots of instantaneous HSN Ca^2+^ transient peak frequencies (and inter-transient intervals) of adult HSN Ca^2+^ activity. Asterisks indicate *p*<0.0001 (Kruskal-Wallis test). (D) Scatterplots show total time spent by each individual with HSN transients ≤30s apart in untreated (green filled circles) and FUDR-treated (green open circles) wild type animals or fertile (blue filled circles) or sterile (blue open circles) *glp-1(or178ts)* mutant animals. Asterisks indicate *p*≤0.0001 (one-way ANOVA); error bars indicate 95% confidence intervals for the mean.

We performed a reciprocal experiment to test how the accumulation of unlaid eggs would affect presynaptic HSN activity. We have previously shown that passage of eggs through the vulva mechanically activates the uv1 neuroendocrine cells which release tyramine and neuropeptides that inhibit HSN activity and egg laying (Collins et al., 2016; Banerjee et al., 2017). We hypothesized that prevention of egg release would block inhibitory uv1 feedback and increase HSN activity. We expressed HisCl channels in the vulval muscles and recorded HSN Ca^2+^ activity after silencing with exogenous histamine. Surprisingly, we found that acute silencing of vulval muscles significantly reduced presynaptic HSN Ca^2+^ activity, resembling the effects of animal sterilization (Fig. 7A and 7B). While untreated animals spent ~16% of recording time with high frequency HSN activity, this was reduced to ~2% of the total recording time in histamine-treated animals (Fig. 7C). These results indicate that postsynaptic vulval muscle activity is required for the burst firing in the presynaptic HSN neurons that accompanies the egg-laying active state.

**Fig. 7.**
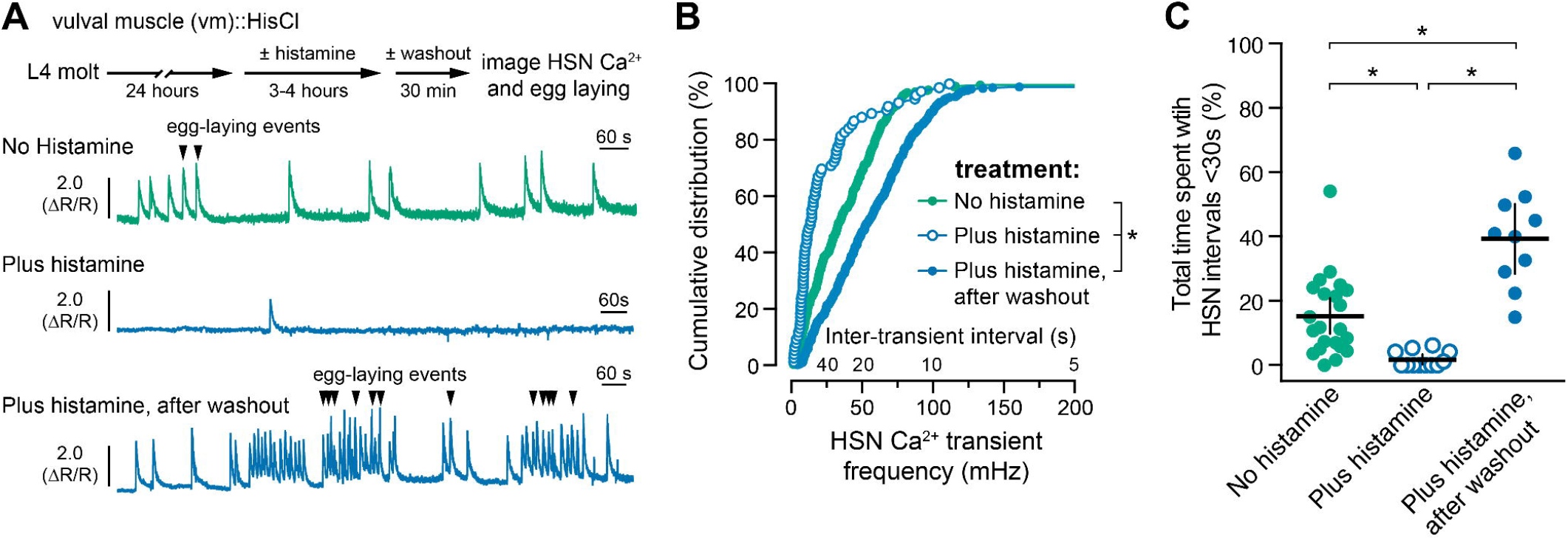
Vulval muscle activity and egg accumulation promote HSN burst firing. (A) 24-hour old adult animals expressing HisCl in the postsynaptic vulval muscles (vm) and GCaMP5/mCherry in the presynaptic HSNs were placed onto NGM plates with (blue, bottom) or without histamine (green, top) for 3-4 hours to induce muscle silencing and cessation of egg laying. Animals were then moved to plates without histamine and allowed to recover for 30 minutes before HSN Ca^2+^ imaging. HSN Ca^2+^ imaging was also performed on adults not removed from histamine (blue, middle). Arrowheads indicate egg laying events. (B) Cumulative distribution plots of instantaneous HSN Ca^2+^ transient peak frequencies (and inter-transient intervals) on histamine (open blue circles), and after histamine washout (filled blue circles) compared with untreated controls (filled green circles). Asterisks indicate *p*<0.0001 (Kruskal-Wallis test). (C) Scatter plots show fraction of time spent by each individual with frequent HSN Ca^2+^ transients characteristic of the egg-laying active state (<30 s) in untreated controls (green circles), on histamine (blue open circles), and after histamine washout (blue circles). Error bars indicate 95% confidence intervals for the mean; asterisks indicate *p*≤0.0061 (one-way ANOVA).

We next looked at how HSN Ca^2+^ activity recovers when histamine inhibition of the vulval muscles and egg laying is reversed. As shown in Fig. 7A, adult animals were treated with or without histamine for 3-4 hours and then moved to plates without histamine for a 20-30 minutes recovery period. Presynaptic HSN Ca^2+^ activity was then recorded as the animals resumed egg-laying behavior. The HSNs showed a rapid and dramatic recovery of Ca^2+^ activity after histamine washout resulting in a prolonged active state with increased HSN Ca^2+^ transient frequency and numerous egg-laying events (Fig. 7A and 7B). Washout animals spent ~40% of their recorded time with elevated HSN activity compared to 15% of the total recorded time in untreated controls (Fig. 7C). During this recovery period, we observed increased vulval muscle twitching contractions in the bright field channel, indicating that muscle activity was restored (data not shown). These results are consistent with a model whereby accumulation of unlaid eggs promotes vulval muscle activity which drives a homeostatic increase in presynaptic HSN activity and burst-firing that sustains egg laying.

HSN synapses are formed exclusively on the lateral vm2 muscle arms that provide sites of contact between the anterior and posterior vulval muscles (White et al., 1986; Feinberg et al., 2008; Collins and Koelle, 2013). Hypomorphic Notch signaling mutants fail to develop vm2 muscle arms, and are egg-laying defective, but have normal pre-synaptic HSN and VC development (Sundaram and Greenwald, 1993; Li et al., 2013). To determine if retrograde signaling from the vulval muscles to the HSNs occurs through the vm2 muscle arms, we recorded HSN Ca^2+^ activity in *lin-12(wy750)* Notch receptor mutant animals (Fig. 8A and 8B). We found that HSN Ca^2+^ transient frequency was strongly reduced in the *lin-12(wy750)* mutants compared to wild-type control animals (Fig. 8C and 8D). HSN Ca^2+^ transients still occurred in *lin-12(wy750)* mutants, but burst-firing was eliminated. Wild-type animals spent ~13% of their time with HSN transients <30 s apart, while this was reduced to zero in the *lin-12(wy750)* mutant (Fig. 8E), resembling activity seen in sterilized or vulval muscle-silenced animals. Together, these results suggest that muscle activity feeds back through the vm2 muscle arms onto the pre-synaptic HSN neurons to promote additional Ca^2+^ transients that drive burst firing and sustain the egg-laying active state.

**Fig. 8:**
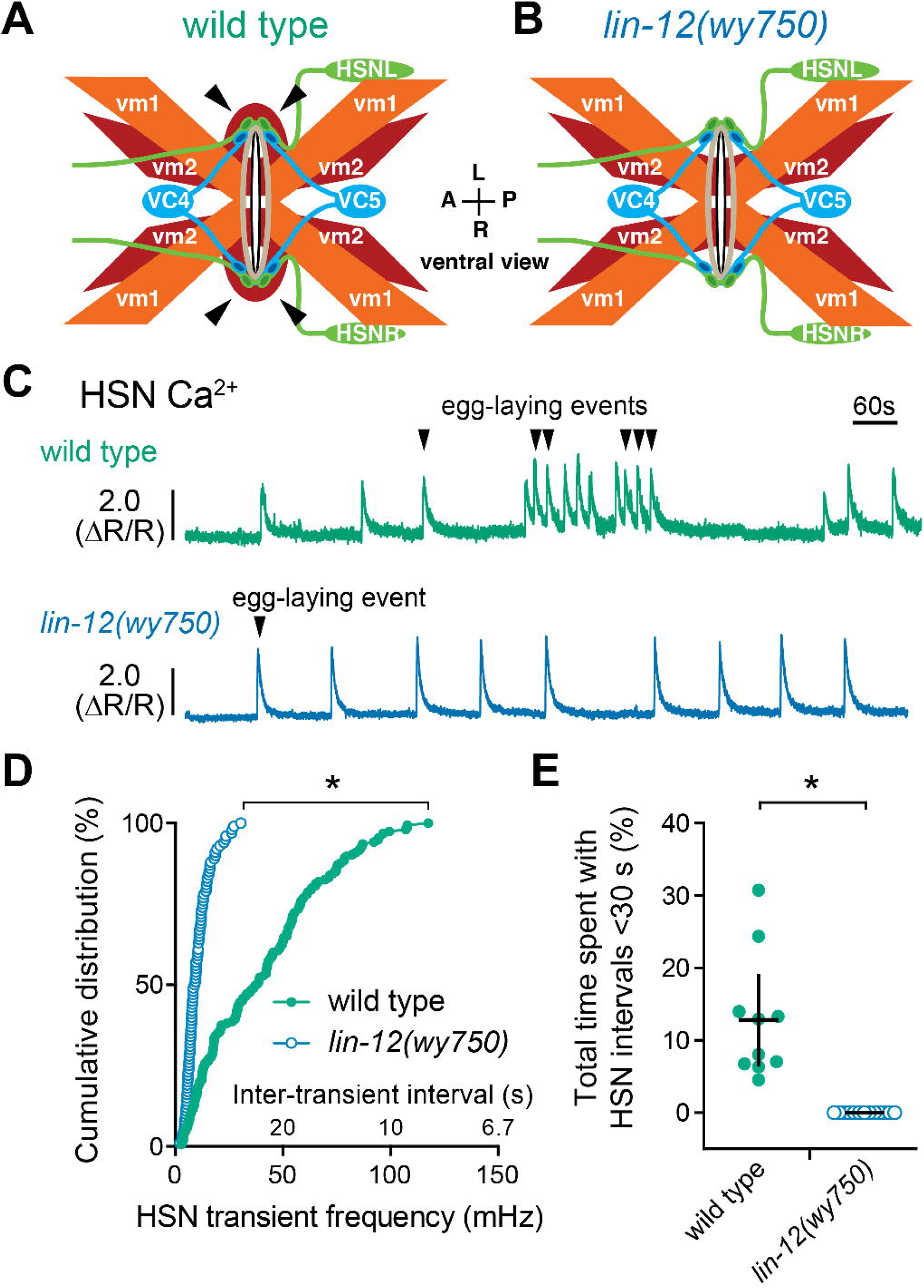
The vm2 muscle arms are required for vulval muscle feedback to HSN and burst firing. (A-B) Cartoon of egg-laying circuit structure (ventral view) in wild-type (A) and *lin-12(wy750)* mutant (B) animals missing lateral vm2 muscle arms (arrowheads). (C) Representative traces show HSN Ca^2+^ activity in wild-type (green) and *lin-12(wy750)* mutant animals (blue). Arrowheads indicate egg-laying events. (D) Cumulative distribution plots of instantaneous Ca^2+^ transient peak frequencies (and inter-transient intervals) in wild-type (green circles) and *lin-12(wy750)* mutants (blue circles). Asterisks indicate *p*<0.0001 (Mann-Whitney test). (E) Scatter plots show fraction of time spent by each individual with frequent HSN Ca^2+^ transients characteristic of the egg-laying active state (<30 s) in wild-type (filled green circles) and *lin-12(wy750)* mutant animals (open blue circles). Error bars indicate 95% confidence intervals for the mean. Asterisk indicates *p*=0.0011 (Student’s t test).

## Discussion

We used a combination of molecular genetic, optogenetic and chemogenetic, and ratiometric Ca^2+^ imaging approaches to determine how coordinated activity develops in the *C. elegans* egg-laying behavior circuit. We find the pre-synaptic HSNs, VCs, and uv1 neuroendocrine cells complete morphological development during early-mid L4 stages, while the vulval muscles finish developing at the late L4 stages. Like HSNs, the vulval muscles show Ca^2+^ activity in the L4.7-8 stage. Coordinated vulval muscle Ca^2+^ transients are not observed until the L4.9 stage when the anterior and posterior vm2 muscle arms complete a Notch-dependent lateral extension around the primary vulval epithelial cells (Li et al., 2013). We do not observe Ca^2+^ transients in the VC neurons and uv1 cells except in egg-laying adults (data not shown) suggesting activity in these cells does not contribute to circuit development. In adults, the juvenile HSN and vulval muscle activity disappears, leading to the establishment of characteristic ‘inactive’ states in which adult animals spend ~85% of their time. Inactive state activity closely resembles that seen in sterilized animals that do not accumulate any eggs. Figure 9 shows a working model for how postsynaptic muscle activity could promote burst firing in the presynaptic HSNs. We propose that uterine cells depress or excite the vulval muscles depending on the degree of stretch. Activation of the uterine muscles, which make gap junctions onto the vm2 muscles, would increase vulval muscle sensitivity to serotonin and other neurotransmitters released from HSN, which subsequently allows for rhythmic acetylcholine input from the VA/VB locomotion motor neurons to drive vulval muscle twitching contractions. Coordinated Ca^2+^ activity in the anterior and posterior vulval muscles diffuses into the vm2 muscle arms to restimulate the HSNs and prolong the egg-laying active state. VC activity is coincident with strong vulval muscle contractions, while uv1 activity follows passage of eggs through the vulva. Once sufficient eggs have been laid, excitatory feedback into the vulval muscles and back to the HSNs is reduced, increasing the probability that inhibitory acetylcholine, tyramine, and neuropeptides released from VC and uv1 will block subsequent HSN Ca^2+^ transients, returning the circuit to the inactive state.

**Fig. 9:**
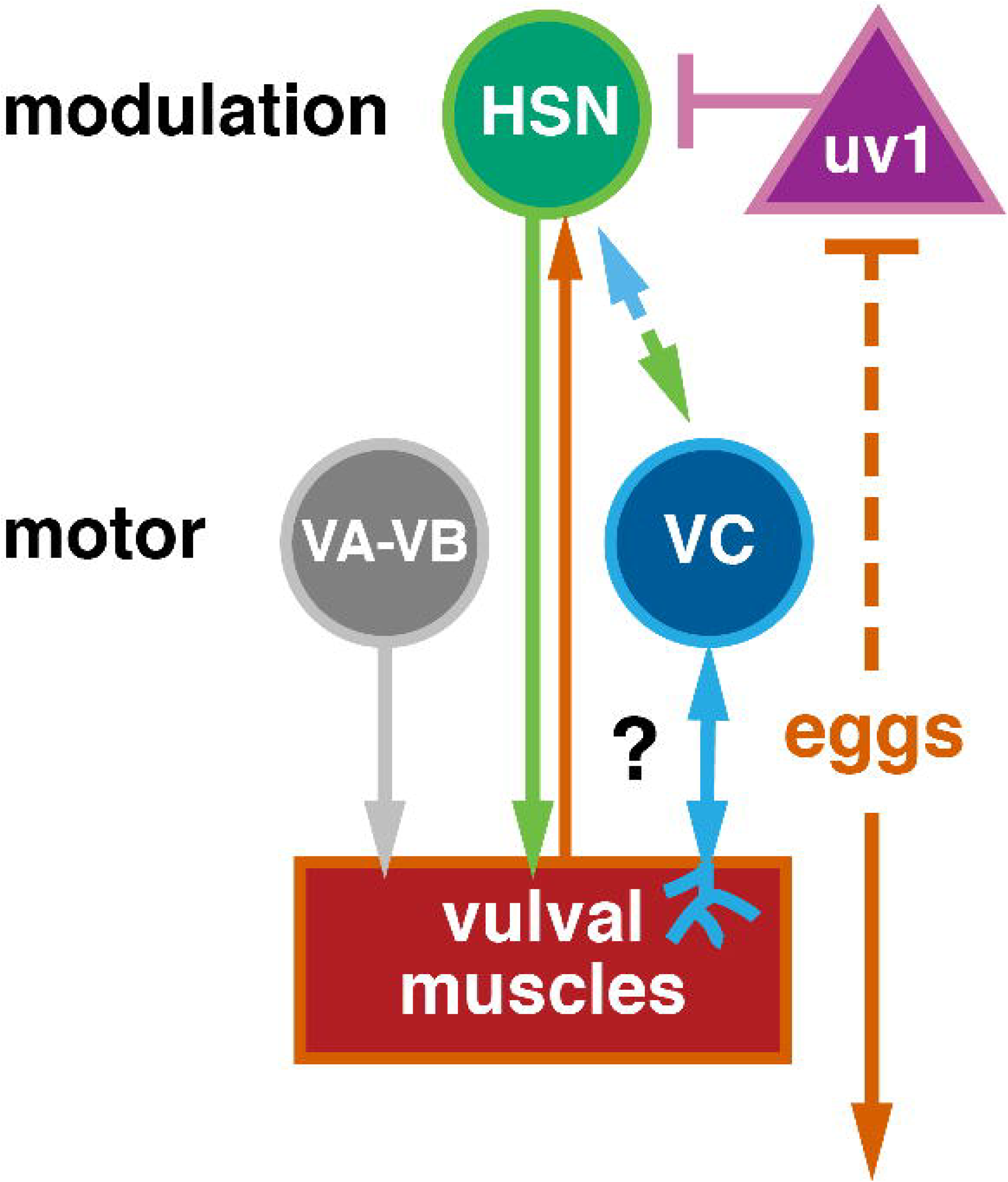
Working model of how retrograde signals from the postsynaptic vulval muscles (question mark) might directly or indirectly modulate burst firing in the presynaptic HSNs. HSN is a serotonergic and peptidergic modulatory command motor neuron that synapses onto the vulval muscles and the VC motor neurons. VA, VB, and VC are cholinergic motor neurons that synapse onto the vulval muscles. uv1 is a tyraminergic and peptidergic neuroendocrine cell mechanically activated by egg release that then feedback inhibits HSN. Arrows indicate activation, bar-headed lines indicate inhibition; see text for more details.

Changes in gene expression likely contribute to the changes in circuit activity patterns we observe between L4s and adults. Previous work has found that serotonin expression is low in L4 and increases as animals increase egg laying (Tanis et al., 2008). Since mutants lacking serotonin have little effect on the timing of the first egg-laying event, we anticipate other neurotransmitters released from the HSNs promote egg laying in young adults. KCC-2 and ABTS-1, two Cl^−^ extruders required for inhibitory neurotransmission, show a developmental increase in HSN expression from L4 to adult (Tanis et al., 2009; Bellemer et al., 2011) which may be associated with the disappearance of spontaneous rhythmic activity in the HSNs after the late L4 stages. At the same time, we find that inhibitory ERG K+ channel expression becomes upregulated in the vulval muscles of young adults. Studies in vertebrate models have shown that mechanical stretch can increase the transcription of receptors that enhance muscle contraction during parturition (Terzidou et al., 2005; Shynlova et al., 2007). We speculate that similar mechanisms may operate in the *C. elegans* reproductive system to drive expression of receptors and channels that modulate vulval muscle sensitivity to presynaptic input. Identifying additional genes whose expression increases upon egg accumulation could help explain how HSN-deficient animals are still able to enter otherwise normal egg-laying active states after sufficient eggs have accumulated.

The HSNs show dramatic changes in Ca^2+^ transient frequency between the inactive and active states. Major G proteins, Gα_q_ and Gα_0_, signal in HSN to increase and inhibit egg laying, respectively (Ringstad and Horvitz, 2008; Tanis et al., 2008). G protein signaling in HSN may modulate an intrinsic pacemaker activity, similar to that seen in other central pattern generator circuits and in the cardiac pacemaker (Hille, 2001). Gα_o_ signaling in HSN activates inhibitory IRK K+ channels (Emtage et al., 2012), and recent work has identified the T-type Ca^2+^ channel, CCA-1, and the Na leak channels, NCA-1 and NCA-2, as possible targets of excitatory Gα_q_ signaling (Yeh et al., 2008; Topalidou et al., 2012; Zang et al., 2017). The balance of both G protein signaling pathways would allow for HSN frequency modulation and dictate whether animals enter or leave the egg-laying active state.

Early vulval muscle activity may be spontaneous or driven by neuronal input. Spontaneous Ca^2+^ transients promote the maturation of activity in many other cells (Moody and Bosma, 2005). We observed no change in behavioral onset or egg-laying rate in animals in which neuron or vulval muscle activity was silenced in the L4 stage. While this may result from incomplete silencing using the HisCl based approach, previous results in other circuits indicate synapse development does not require Ca^2^+-dependent excitatory transmission (Verhage et al., 2000; Lu et al., 2013; Sando et al., 2017). While G protein signaling may drive early Ca^2+^ activity in the absence of electrical activity, synaptic transmission would still require Ca^2^+-dependent vesicle fusion. The persistence of vulval muscle activity in animals that lack HSNs and its recovery after acute neural silencing suggests the activity we observe arises from a shared mechanism that is not strictly required for synapse development and/or recovers quickly after histamine washout.

Our work continues to show the functional importance of the post-synaptic vm2 muscle arms in coordinating muscle activity during egg-laying behavior. Because of the intervening vulval slit through which eggs are laid, the vm2 muscle arms are the only sites of contact between the anterior and posterior muscles. Coordinated muscle Ca^2+^ transients appear during the L4.9 larval stage after vm2 muscle arm development. After development, the vm2 muscle arms may be electrically coupled at their points of contact, allowing for the immediate spread of electrical activity and/or Ca^2+^ signals between the anterior and posterior muscles. In addition to uncoordinated vm1 and vm2 Ca^2+^ activity, mutants missing the vm2 muscle arms do not show regenerative HSN Ca^2+^ activity, resembling the consequences of vulval muscle electrical silencing (Li et al., 2013). The vm2 muscle arms also form the sites of synaptic input from HSN and VC. We have previously shown that the ERG K+ channel and SER-1 serotonin receptor localize to the vm2 muscle arm region (Collins and Koelle, 2013; Li et al., 2013). Both ERG and SER-1 have C-terminal PDZ interaction motifs, and SER-1 has been shown to interact with the large PDZ scaffold protein MPZ-1 that may drive the local organization of these and other molecules to the vm2 muscle arms (Xiao et al., 2006). Innexin gap junction proteins which are potential targets of G protein signaling (Correa et al., 2015) may also play a role in driving the development of coordinated vulval muscle contractility and HSN ‘burst’ activity in the circuit during egg laying.

The importance of stretch-mediated feedback is well characterized in circuits that control autonomic functions (Dethier and Gelperin, 1967; Gelperin, 1971; Spencer et al., 2002), the rhythmic uterine activity during parturition (Ferguson’s reflex) (Ferguson, 1941), and in circuits which generate rhythmic motor outputs (Grillner, 2003; Marder et al., 2005; Blitz and Nusbaum, 2011). Stretch can provide either positive or negative feedback to downstream reflex and homeostatic circuits. For example, specialized mechanosensory neurons activated by gastric stretch induce satiety by providing negative feedback to neural circuits controlling food consumption (Dethier and Gelperin, 1967; Zagorodnyuk et al., 2001). In guinea pigs, stretch-sensitive interneurons provide ascending excitatory and descending inhibitory inputs to generate peristaltic neural reflexes in the distal colon (Spencer and Smith, 2004). Mechanical stretch (from egg accumulation) or artificially induced distension of the reproductive tract in female flies induces an attraction to acetic acid so that eggs can be laid in optimal environments (Gou et al., 2014). In the cases described above, how stretch sensory inputs modulate the activity of neural circuits and synaptic transmission is not always clear.

The *C. elegans* egg-laying homeostat is regulated by egg accumulation which sustains rhythmic activity in a motor neuron for muscle contraction and egg release. In the case of the Ferguson’s parturition reflex, initial stretch-induced myogenic contractions engage the neuroendocrine feed-forward loop, similar to our results showing that vulval muscle activity promotes a feed-forward increase in HSN activity. Does mechanosensory stretch also play a role in the feedback inhibition of *C. elegans* egg-laying? While the release of eggs and loss of uterine stretch should decrease feedforward drive into the vulval muscles and HSN, additional mechanical feedback from the VC motor neurons and the uv1 neuroendocrine cells may be required to exit the active state completely. VC Ca^2+^ activity is coincident with egg release, and mutants with reduced acetylcholine or VC function have more frequent egg-laying events (Bany et al., 2003). The uv1 cells are mechanically deformed and activated by egg release, and tyramine and inhibitory neuropeptides released from uv1 inhibit HSN activity (Collins et al., 2016; Banerjee et al., 2017). Further studies of the *C. elegans* egg-laying homeostat described here should allow the dissection of conserved molecular, cellular, and synaptic mechanisms that drive stretch-dependent feedback.

## Movie legends

Movie 1. Ratio recording of a HSN Ca^2+^ transient at the L4.9 larval stage. High Ca^2+^ is indicated in red while low calcium is in blue. The HSN cell body and pre-synaptic terminal are indicated. Head is at bottom, tail is at left.

Movie 2. Ratio recording of a HSN Ca^2+^ transient prior to an egg-laying event in an adult animal during the active state. High Ca^2+^ is indicated in red while low calcium is in blue. The HSN cell body and pre-synaptic terminal are indicated. Head is at bottom, tail is at top.

Movie 3. Ratio recording of an uncoordinated vulval muscle Ca^2+^ transient at the L4.7-8 larval stage. High Ca^2+^ is indicated in red while low calcium is in blue. Developing anterior and posterior vulval muscles are indicated. Head is at top, tail is at bottom.

Movie 4. Ratio recording of an uncoordinated vulval muscle Ca^2+^ transient at the L4.9 larval stage. High Ca^2+^ is indicated in red while low calcium is in blue. Anterior and posterior vulval muscles are indicated. Head is at left, tail is at bottom.

Movie 5. Ratio recording of a coordinated vulval muscle Ca^2+^ transient at the L4.9 larval stage. High Ca^2+^ is indicated in red while low calcium is in blue. Anterior and posterior vulval muscles are indicated. Head is at top, tail is at bottom.

Movie 6. Ratio recording of coordinated vulval muscle Ca^2+^ transients during egg laying in adult animals. High Ca^2+^ is indicated in red while low calcium is in blue. The anterior and posterior vulval muscles are indicated along with a previously laid egg. Head is at right, tail is at left.

## Acknowledgements

This work was funded by a grant from NINDS to KMC (R01 NS086932). JG was supported by the Bridge to the Baccalaureate Program (R25 GM050083). Strains used in this study have been provided to the *C. elegans* Genetics Center, which is funded by NIH Office of Research Infrastructure Programs (P40 OD010440). We thank Addys Bode and Michael Scheetz for technical assistance and help with strain construction. We thank James Baker, Julia Dallman, Laura Bianchi, Peter Larsson, Stephen Roper, and members of the Collins lab for helpful discussions and feedback.

## References

Adler CE, Fetter RD, Bargmann CI (2006) UNC-6/Netrin induces neuronal asymmetry and defines the site of axon formation. Nat Neurosci 9:511–518.

Akerboom J et al. (2013) Genetically encoded calcium indicators for multi-color neural activity imaging and combination with optogenetics. Front Mol Neurosci 6:2.

Banerjee N, Bhattacharya R, Gorczyca M, Collins KM, Francis MM (2017) Local neuropeptide signaling modulates serotonergic transmission to shape the temporal organization of *C. elegans* egg-laying behavior. PLoS Genet 13:e1006697.

Bany IA, Dong MQ, Koelle MR (2003) Genetic and cellular basis for acetylcholine inhibition of *Caenorhabditis elegans* egg-laying behavior. J Neurosci 23:8060–8069.

Bellemer A, Hirata T, Romero MF, Koelle MR (2011) Two types of chloride transporters are required for GABA(A) receptor-mediated inhibition in *C. elegans*. EMBO J 30:1852–1863.

Blitz DM, Nusbaum MP (2011) Neural circuit flexibility in a small sensorimotor system. Curr Opin Neurobiol 21:544–552.

Borodinsky LN, Root CM, Cronin JA, Sann SB, Gu X, Spitzer NC (2004) Activity-dependent homeostatic specification of transmitter expression in embryonic neurons. Nature 429:523–530.

Brenner S (1974) The genetics of *Caenorhabditis elegans*. Genetics 77:71–94.

Cai T, Fukushige T, Notkins AL, Krause M (2004) Insulinoma-Associated Protein IA-2, a Vesicle Transmembrane Protein, Genetically Interacts with UNC-31/CAPS and Affects Neurosecretion in *Caenorhabditis elegans*. J Neurosci 24:3115–3124.

Carnell L, Illi J, Hong SW, McIntire SL (2005) The G-protein-coupled serotonin receptor SER-1 regulates egg laying and male mating behaviors in *Caenorhabditis elegans*. J Neurosci 25:10671–10681.

Chase DL, Pepper JS, Koelle MR (2004) Mechanism of extrasynaptic dopamine signaling in *Caenorhabditis elegans*. Nat Neurosci 7:1096–1103.

Clark SG, Lu X, Horvitz HR (1994) The *Caenorhabditis elegans* locus lin-15, a negative regulator of a tyrosine kinase signaling pathway, encodes two different proteins. Genetics 137:987–997.

Colavita A, Tessier-Lavigne M (2003) A Neurexin-related protein, BAM-2, terminates axonal branches in *C. elegans*. Science 302:293–296.

Collins KM, Koelle MR (2013) Postsynaptic ERG potassium channels limit muscle excitability to allow distinct egg-laying behavior states in *Caenorhabditis elegans*. J Neurosci 33:761–775.

Collins KM, Bode A, Fernandez RW, Tanis JE, Brewer JC, Creamer MS, Koelle MR (2016) Activity of the *C. elegans* egg-laying behavior circuit is controlled by competing activation and feedback inhibition. Elife 5:e21126.

Correa PA, Gruninger T, Garcia LR (2015) DOP-2 D2-Like Receptor Regulates UNC-7 Innexins to Attenuate Recurrent Sensory Motor Neurons during *C. elegans* Copulation. J Neurosci 35:9990–10004.

Dempsey CM, Mackenzie SM, Gargus A, Blanco G, Sze JY (2005) Serotonin (5HT), fluoxetine, imipramine and dopamine target distinct 5HT receptor signaling to modulate *Caenorhabditis elegans* egg-laying behavior. Genetics 169:1425–1436.

Dernovici S, Starc T, Dent JA, Ribeiro P (2007) The serotonin receptor SER-1 (5HT2ce) contributes to the regulation of locomotion in *Caenorhabditis elegans*. Dev Neurobiol 67:189–204.

Dethier VG, Gelperin A (1967) Hyperphagia in the Blowfly. Journal of Experimental Biology 47:191–200.

Emtage L, Aziz-Zaman S, Padovan-Merhar O, Horvitz HR, Fang-Yen C, Ringstad N (2012) IRK-1 potassium channels mediate peptidergic inhibition of *Caenorhabditis elegans* serotonin neurons via a G(o) signaling pathway. J Neurosci 32:16285–16295.

Feinberg EH, Vanhoven MK, Bendesky A, Wang G, Fetter RD, Shen K, Bargmann CI (2008) GFP Reconstitution Across Synaptic Partners (GRASP) defines cell contacts and synapses in living nervous systems. Neuron 57:353–363.

Ferguson JKW (1941) A study of the motility of the intact uterus at term. Surg Gynaecol Obste:359–366.

Flavell SW, Pokala N, Macosko EZ, Albrecht DR, Larsch J, Bargmann CI (2013) Serotonin and the neuropeptide PDF initiate and extend opposing behavioral states in *C. elegans*. Cell 154:1023–1035.

Fujiwara M, Aoyama I, Hino T, Teramoto T, Ishihara T (2016) Gonadal Maturation Changes Chemotaxis Behavior and Neural Processing in the Olfactory Circuit of *Caenorhabditis elegans*. Curr Biol 26:1522–1531.

Garaschuk O, Hanse E, Konnerth A (1998) Developmental profile and synaptic origin of early network oscillations in the CA1 region of rat neonatal hippocampus. J Physiol 507 (Pt 1):219–236.

Garaschuk O, Linn J, Eilers J, Konnerth A (2000) Large-scale oscillatory calcium waves in the immature cortex. Nat Neurosci 3:452–459.

Gelperin A (1971) Abdominal sensory neurons providing negative feedback to the feeding behavior of the blowfly. Zeitschrift für vergleichende Physiologie 72:17–31.

Gou B, Liu Y, Guntur AR, Stern U, Yang CH (2014) Mechanosensitive neurons on the internal reproductive tract contribute to egg-laying-induced acetic acid attraction in Drosophila. Cell Rep 9:522–530.

Grillner S (2003) The motor infrastructure: from ion channels to neuronal networks. Nat Rev Neurosci 4:573–586.

Gu X, Spitzer NC (1995) Distinct aspects of neuronal differentiation encoded by frequency of spontaneous Ca^2+^ transients. Nature 375:784–787.

Gu X, Olson EC, Spitzer NC (1994) Spontaneous neuronal calcium spikes and waves during early differentiation. J Neurosci 14:6325–6335.

Gurel G, Gustafson MA, Pepper JS, Horvitz HR, Koelle MR (2012) Receptors and other signaling proteins required for serotonin control of locomotion in *Caenorhabditis elegans*. Genetics 192:1359–1371.

Hanson MG, Milner LD, Landmesser LT (2008) Spontaneous rhythmic activity in early chick spinal cord influences distinct motor axon pathfinding decisions. Brain Res Rev 57:77–85.

Hapiak VM, Hobson RJ, Hughes L, Smith K, Harris G, Condon C, Komuniecki P, Komuniecki RW (2009) Dual excitatory and inhibitory serotonergic inputs modulate egg laying in *Caenorhabditis elegans*. Genetics 181:153–163.

Hardaker LA, Singer E, Kerr R, Zhou G, Schafer WR (2001) Serotonin modulates locomotory behavior and coordinates egg-laying and movement in *Caenorhabditis elegans*. J Neurobiol 49:303–313.

Harfe BD, Fire A (1998) Muscle and nerve-specific regulation of a novel NK-2 class homeodomain factor in *Caenorhabditis elegans*. Development 125:421–429.

Hille B (2001) Ion Channels of Excitable Membranes, Third Edition: Sinaur.

Hobson RJ, Hapiak VM, Xiao H, Buehrer KL, Komuniecki PR, Komuniecki RW (2006) SER-7, a *Caenorhabditis elegans* 5-HT7-like receptor, is essential for the 5-HT stimulation of pharyngeal pumping and egg laying. Genetics 172:159–169.

Jarecki J, Keshishian H (1995) Role of neural activity during synaptogenesis in *Drosophila*. J Neurosci 15:8177–8190.

Li C, Chalfie M (1990) Organogenesis in *C. elegans*: positioning of neurons and muscles in the egg-laying system. Neuron 4:681–695.

Li P, Collins KM, Koelle MR, Shen K (2013) LIN-12/Notch signaling instructs postsynaptic muscle arm development by regulating UNC-40/DCC and MADD-2 in *Caenorhabditis elegans*. Elife 2:e00378.

Lu W, Bushong EA, Shih TP, Ellisman MH, Nicoll RA (2013) The cell-autonomous role of excitatory synaptic transmission in the regulation of neuronal structure and function. Neuron 78:433–439.

Marder E, Bucher D, Schulz DJ, Taylor AL (2005) Invertebrate central pattern generation moves along. Curr Biol 15:R685–699.

Mok DZ, Sternberg PW, Inoue T (2015) Morphologically defined sub-stages of *C. elegans* vulval development in the fourth larval stage. BMC Dev Biol 15:26.

Moody WJ, Bosma MM (2005) Ion channel development, spontaneous activity, and activity-dependent development in nerve and muscle cells. Physiol Rev 85:883–941.

Moresco JJ, Koelle MR (2004) Activation of EGL-47, a Galpha(o)-coupled receptor, inhibits function of hermaphrodite-specific motor neurons to regulate *Caenorhabditis elegans* egg-laying behavior. J Neurosci 24:8522–8530.

Patel MR, Lehrman EK, Poon VY, Crump JG, Zhen M, Bargmann CI, Shen K (2006) Hierarchical assembly of presynaptic components in defined *C. elegans* synapses. Nat Neurosci 9:1488–1498.

Pokala N, Liu Q, Gordus A, Bargmann CI (2014) Inducible and titratable silencing of *Caenorhabditis elegans* neurons in vivo with histamine-gated chloride channels. Proc Natl Acad Sci U S A 111:2770–2775.

Raizen DM, Zimmerman JE, Maycock MH, Ta UD, You YJ, Sundaram MV, Pack AI (2008) Lethargus is a *Caenorhabditis elegans* sleep-like state. Nature 451:569–572.

Ravi B, Nassar LM, Kopchock III RJ, Dhakal P, Scheetz M, Collins KM (2018) Ratiometric calcium imaging of individual neurons in behaving *Caerhabditis elegans*. J Vis Exp:e56911.

Ringstad N, Horvitz HR (2008) FMRFamide neuropeptides and acetylcholine synergistically inhibit egg-laying by *C. elegans*. Nat Neurosci 11:1168–1176.

Sando R, Bushong E, Zhu Y, Huang M, Considine C, Phan S, Ju S, Uytiepo M, Ellisman M, Maximov A (2017) Assembly of Excitatory Synapses in the Absence of Glutamatergic Neurotransmission. Neuron 94:312–321 e313.

Schindelin J, Arganda-Carreras I, Frise E, Kaynig V, Longair M, Pietzsch T, Preibisch S, Rueden C, Saalfeld S, Schmid B, Tinevez JY, White DJ, Hartenstein V, Eliceiri K, Tomancak P, Cardona A (2012) Fiji: an open-source platform for biological-image analysis. Nat Methods 9:676–682.

Shen K, Bargmann CI (2003) The immunoglobulin superfamily protein SYG-1 determines the location of specific synapses in *C. elegans*. Cell 112:619–630.

Shen K, Fetter RD, Bargmann CI (2004) Synaptic specificity is generated by the synaptic guidepost protein SYG-2 and its receptor, SYG-1. Cell 116:869–881.

Shynlova O, Williams SJ, Draper H, White BG, MacPhee DJ, Lye SJ (2007) Uterine stretch regulates temporal and spatial expression of fibronectin protein and its alpha 5 integrin receptor in myometrium of unilaterally pregnant rats. Biol Reprod 77:880–888.

Spencer NJ, Smith TK (2004) Mechanosensory S-neurons rather than AH-neurons appear to generate a rhythmic motor pattern in guinea-pig distal colon. J Physiol 558:577–596.

Spencer NJ, Hennig GW, Smith TK (2002) A rhythmic motor pattern activated by circumferential stretch in guinea-pig distal colon. J Physiol 545:629–648.

Sundaram M, Greenwald I (1993) Suppressors of a lin-12 hypomorph define genes that interact with both lin-12 and glp-1 in *Caenorhabditis elegans*. Genetics 135:765–783.

Tanis JE, Moresco JJ, Lindquist RA, Koelle MR (2008) Regulation of serotonin biosynthesis by the G proteins Galphao and Galphaq controls serotonin signaling in *Caenorhabditis elegans*. Genetics 178:157–169.

Tanis JE, Bellemer A, Moresco JJ, Forbush B, Koelle MR (2009) The potassium chloride cotransporter KCC-2 coordinates development of inhibitory neurotransmission and synapse structure in *Caenorhabditis elegans*. J Neurosci 29:9943–9954.

Terzidou V, Sooranna SR, Kim LU, Thornton S, Bennett PR, Johnson MR (2005) Mechanical stretch up-regulates the human oxytocin receptor in primary human uterine myocytes. J Clin Endocrinol Metab 90:237–246.

Topalidou I, Keller C, Kalebic N, Nguyen KC, Somhegyi H, Politi KA, Heppenstall P, Hall DH, Chalfie M (2012) Genetically separable functions of the MEC-17 tubulin acetyltransferase affect microtubule organization. Curr Biol 22:1057–1065.

Tsalik EL, Hobert O (2003) Functional mapping of neurons that control locomotory behavior in *Caenorhabditis elegans*. J Neurobiol 56:178–197.

Verhage M, Maia AS, Plomp JJ, Brussaard AB, Heeroma JH, Vermeer H, Toonen RF, Hammer RE, van den Berg TK, Missler M, Geuze HJ, Sudhof TC (2000) Synaptic assembly of the brain in the absence of neurotransmitter secretion. Science 287:864–869.

Waggoner LE, Zhou GT, Schafer RW, Schafer WR (1998) Control of alternative behavioral states by serotonin in *Caenorhabditis elegans*. Neuron 21:203–214.

Warp E, Agarwal G, Wyart C, Friedmann D, Oldfield CS, Conner A, Del Bene F, Arrenberg AB, Baier H, Isacoff EY (2012) Emergence of patterned activity in the developing zebrafish spinal cord. Curr Biol 22:93–102.

Watt AJ, Cuntz H, Mori M, Nusser Z, Sjostrom PJ, Hausser M (2009) Traveling waves in developing cerebellar cortex mediated by asymmetrical Purkinje cell connectivity. Nat Neurosci 12:463–473.

White JG, Southgate E, Thomson JN, Brenner S (1986) The structure of the nervous system of the nematode *Caenorhabditis elegans*. Philos Trans R Soc Lond B Biol Sci 314:1–340.

Wong RO, Chernjavsky A, Smith SJ, Shatz CJ (1995) Early functional neural networks in the developing retina. Nature 374:716–718.

Xiao H, Hapiak VM, Smith KA, Lin L, Hobson RJ, Plenefisch J, Komuniecki R (2006) SER-1, a *Caenorhabditis elegans* 5-HT2-like receptor, and a multi-PDZ domain containing protein (MPZ-1) interact in vulval muscle to facilitate serotonin-stimulated egg-laying. Dev Biol 298:379–391.

Yeh E, Ng S, Zhang M, Bouhours M, Wang Y, Wang M, Hung W, Aoyagi K, Melnik-Martinez K, Li M, Liu F, Schafer WR, Zhen M (2008) A putative cation channel, NCA-1, and a novel protein, UNC-80, transmit neuronal activity in *C. elegans*. PLoS Biol 6:e55.

Zagorodnyuk VP, Chen BN, Brookes SJ (2001) Intraganglionic laminar endings are mechano-transduction sites of vagal tension receptors in the guinea-pig stomach. J Physiol 534:255–268.

Zang KE, Ho E, Ringstad N (2017) Inhibitory peptidergic modulation of *C. elegans* serotonin neurons is gated by T-type calcium channels. Elife 6:e22771.

Zhang M, Chung SH, Fang-Yen C, Craig C, Kerr RA, Suzuki H, Samuel AD, Mazur E, Schafer WR (2008) A self-regulating feed-forward circuit controlling C. elegans egg-laying behavior. Curr Biol 18:1445–1455.

